# Identification and validation of E3 ubiquitin ligase XIAP as a novel substrate of deubiquitinase USP7 (HAUSP) - Implication towards oncogenesis

**DOI:** 10.1101/2021.08.12.456108

**Authors:** Gouranga Saha, Sibani Sarkar, Partha S Mohanta, Krishna Kumar, Saikat Chakraborty, Mrinal K Ghosh

## Abstract

The induction of apoptosis upon USP7 (HAUSP) inhibition is established in cancers that contain a wild-type p53 (p53^Wt^) through the ‘USP7-Mdm2-p53’ axis, but no clear explanation has yet been reported for the same to occur in cancers containing mutant 53 (p53^Mut^) or even p53 null (p53^Null^) systems. Instead of this ‘USP7-Mdm2-p53’ axis USP7 also works through an alternative new pathway identified in this study. Here in this study, we observed that the magnitude of apoptosis induction in response to USP7 inhibition was remarkably similar between cancer cells showing p53^Null^ or p53^Mut^ and those with p53^Wt^. Through a proteomics-based approach, we were able to identify XIAP as a novel interacting partner for USP7. XIAP is a potent and well-characterized member of the inhibitor of apoptosis proteins (IAPs), which function through caspase inhibition. We successfully identified USP7 as a positive regulator of XIAP at post-translational but not at its transcriptional level. Using molecular modelling coupled with domain deletion studies, we show that the first three Ubl domains in association with the catalytic domain of USP7 interact with the BIR2 and the linker region between BIR2 and BIR3 domains of XIAP. Modulation of expression and catalytic activity of USP7 in multiple type of cancer cell lines showed that USP7 stabilizes XIAP through its deubiquitinase activity. We have also observed that USP7 sensitizes cells against chemotherapeutic drugs through stabilization of XIAP. Thus, USP7 promotes tumorigenesis in multiple cancers, *via* stabilization of XIAP that results in apoptosis inhibition in caspase dependent pathway. Moreover, we observed that combinatorial inhibition of USP7 and XIAP can induce cellular apoptosis in a higher magnitude than their individual inhibition. Additionally, our results indicates that nanoformulated P5091 and P22077 showed higher potency for killing C6 cells in comparison to normal drugs. To the best of our knowledge, this is the first report on identification and validation of XIAP, a crucial E3 ubiquitin ligase, as a novel substrate of the deubiquitinase USP7 and they together involve in empowerment of the tumorigenic potential of cancer cells.

## Introduction

Programmed cell death or apoptosis is an evolutionary conserved and highly regulated process that utilized by multicellular organisms during development, immune responses, and ageing. Apoptosis may occur by either extrinsic or intrinsic modes of activation. The primary goal of both the mechanisms is the activation of caspase-cascade, which sets up a series of events known as the execution phase which ultimately destroys the resident cells^1^. A class of proteins known as inhibitor of apoptosis proteins (IAPs) functions as negative regulators of caspases. At present, there are eight known mammalian IAPs – *viz.,* neuronal IAP (NAIP), cellular IAP1 (cIAP1), cellular IAP2 (cIAP2), X-linked IAP (XIAP), Survivin, BIR-containing ubiquitin-conjugating enzyme (BRUCE), melanoma IAP (ML-IAP), and IAP-like protein 2 (ILP2)^2^. IAPs are frequently over-expressed in various human cancers, thereby contributing to tumor survival, progression, and chemoresistance. As a result, IAPs have been targeted in many anticancer therapies^3^. XIAP is the most characterized member of all IAPs and has also been argued to be the most potent IAP in terms of anti-apoptotic ability. It has three baculovirus IAP repeat (BIR) domains by which it interacts with target proteins^4^. The BIR3 domain allows XIAP to interact with caspase-9, whereas BIR2 and the linker region between BIR2 and BIR1 facilitate the interaction with caspase-3 and 7^5, 6^. The C-terminal portion of XIAP contains a ubiquitin association (UBA) domain and a RING E3 ligase domain used in substrate ubiquitination and targeting them for proteasome-mediated degradation^4^. XIAP has been shown to exhibit aberrant expression patterns throughout a broad spectrum of human cancers. Its overexpression has been linked to chemoresistance and overall poor prognosis in specific patient subgroups^7–11^.

The Ubiquitin Specific Protease 7 (USP7/HAUSP) is a deubiquitinase that modifies the length of poly-ubiquitin chains on its target proteins and has been shown to fulfil different roles in various biological processes ranging from viral infections to malignant transformation^12^. The well-known substrate of USP7 is the p53-MDM2 complex, and several studies have demonstrated that the induction of apoptosis upon USP7 functional inhibition is primarily due to the restoration of p53^13^. Regulation of the p53 pathway by USP7 is one of the well-characterized and heavily studied field of USP7 function. Activated p53 transcriptionally regulates multiple genes involved in several biological processes, including DNA damage repair, cell cycle arrest, apoptosis and senescence. In the context of DNA damage, p53 selectively regulates the expression of multiple genes, which made p53 a decision-making transcription factor in determining cellular fate. In p53^Wt^ containing cancers, the induction of apoptosis by USP7 inhibition thought to work primarily through the ‘USP7-MDM2-p53’ axis. Apart from this classical route of apoptosis induction through the ‘USP7-MDM2-p53’ axis, we have observed that the USP7 small molecule inhibitor P5091 can induce apoptosis in cancer cells irrespective of cellular p53 status. However, the mode of action in p53^Mut^ or p53^Null^ cancers is still a matter of investigation.

To answer this question, we adopted a proteomics-based approach to screen novel cancer-associated USP7 interactors, wherein we successfully identified the anti-apoptotic protein XIAP to be a major candidate. We also analyzed the change in proteome upon USP7 inhibition in both p53^Wt^ and p53^Null^ HCT116 cells to delaminate the pathways functioning under USP7 to control programmed cell death. We then validated and established the nature of USP7-XIAP interaction and its consequential effects in human embryonic kidney (HEK) and different cancer cell lines. Here, in this report, we found USP7 physically interacts with XIAP and stabilizes it by deubiquitination in order to curb apoptosis, augment tumor survival and sensitizing cells towards genotoxic drug-induced apoptotic cell death in a p53-independent manner.

Additionally, due to the diverse role of USP7 in promoting oncogenesis, it should be an ideal target for therapeutic intervention. Currently, several synthetic small molecules have been established as potential inhibitors of USP7 and some are more effective in cancer treatment. USP7 inhibitors developed till date are less bioavailable because of their hydrophobic nature and rapid clearance from circulation, which is one of the major concern. To overcome these problems nanotechnology based approach will be an ideal strategy to enhance the half-life, targeting efficiencies and to achieve the minimum side effects of the molecules. Hence, FDA approved Poly (lactic-co-glycolic acid:PLGA), a biocompatible polymer, has been used to prepare the nanoformulations of USP7 inhibitors for therapeutic application.

## Material and methods

### Antibodies and reagents

The following reagents were purchased from the designated suppliers: Embelin, MG-132, Cycloheximide, Etoposide, and Doxorubicin from Sigma. The antibodies against the following proteins were purchased from the indicated sources: XIAP, PARP, GFP, Caspase7, Cleaved Caspase7, Caspase3, Cleaved Caspase3 Caspase9, Cleaved Caspase9, and Cleaved PARP from Cell Signalling Technology); p53, p21, Ub, and GAPDH from Santa Cruz Biotechnology; Mdm2, Flag, Actin and all HRP-conjugated secondary antibodies from Sigma.

### Cell culture and transfection

Human colorectal cancer (CRC) cell lines [HCT 116 (p53^Wt^ and p53^Null^), SW-480, HT-29, SW-620]; human embryonic kidney cell line [HEK-293 and HEK-293T] and other cell lines (RAW, C6, U87, HeLa, SiHa, HepG2, Huh7) were used (ATCC. Manassas, VA, USA). The cell lines were cultured in McCoy’s 5A, DMEM or RPMI-1640 supplemented with 10% heat-inactivated foetal bovine serum (FBS) and maintained at 5% CO2, 37 °C in a humid incubator as per standard cultured condition specified by ATCC. In addition, Penicillin/Streptomycin cocktail and Gentamycin (Invitrogen, Life Technologies) were added as prescribed by the manufacturer. Transfections of different DNA constructs were performed using Lipofectamine 2000 and 3000 (Invitrogen). HEK293 and HEK293T cell lines were transfected by the Calcium-Phosphate method as described earlier^14^. Lipofectamine RNAi MAX (Invitrogen) was used for siRNA-mediated knockdown experiments.

### Plasmids, shRNA, and siRNA

Small interfering RNAs (siRNAs) against USP7 were purchased from Santa Cruz Biotechnology (Santa Cruz, CA, USA). Three independent shRNAs against USP7 in *pmko.1* based vector was cloned in the lab. XIAP was subcloned from pGEX-XIAP (Addgene Plasmid # 8340) in pEGFP-c3, pIRES-hrGFP-1a and pCDNA4. USP7 expression vector pCI-neo-Flag backbone (Addgene Plasmid # 16655). All the constructs were verified by restriction digestion and confirmed by sequencing. All primers were synthesized from IDT (Integrated DNA Technologies, Coralville, IA, USA). Sequences of the primers are given in Additional file 1.

### Immunoprecipitation (IP)

Cells were resuspended in IP-Lysis buffer (50mM Tris-Cl pH 8, 150 mM NaCl, 0.5% Triton X-100, 0.5% NP-40, 0.5% deoxycholate, 1mM PMSF, 1mM DTT, 10% glycerol) supplemented with protease inhibitor cocktail (Calbiochem). Suspension was kept on ice for 2hr, and then precleared with Protein-A/G Sepharose beads for an additional 2 hrs with gentle rotation at 4^0^C. Precleared lysate was incubated with antibodies of interest overnight at 4^0^C. Next morning the mixture was incubated with blocked beads for 4 hrs at RT. Beads were washed thrice with IP-Lysis buffer and boiled in 2X loading buffer to elute proteins before running SDS PAGE followed by Immunoblotting (IB). For Input, 3% lysate was run separately and probed with indicated antibodies.

### Immunoblot (IB) analysis

Sample for IB was prepared as per the protocol described earlier with some modification^15^. Cells were resuspended in whole-cell lysis buffer and sonicated (30% Amp, 10 sec ON/OFF cycle for 30 sec) then centrifuged at 12000g for 15 min. Each sample (60ug) was boiled in 2X loading buffer and resolved in SDS-PAGE and transferred onto PVDF membrane. Membranes were blocked in 5% BSA in TBS-T and incubated with indicated antibodies diluted in 2.5% BSA in TBS-T overnight at 4^0^C. After incubation with secondary antibody for 2 hrs at room temperature, the membrane was developed using Classico ECL chemiluminescence substrate, Millipore. Images were captured using Bio-Rad Chemi-doc. During capture, a nearly identical standard setting for brightness and contrast was maintained.

### GST-tagged proteins purification and pulldown assay

GST-USP7, GST-XIAP, and GST-Control vectors were transformed and expressed in BL21DE3 cells to purify recombinant proteins. Freshly grown cultures were induced with (1 mmol/l) IPTG at 16°C for overnight. The cells were then harvested, and the proteins purified, concentrated, and performed pulldown experiments following our previous protocols ^16, 17^. The purity of the proteins was checked through SDS–PAGE followed by Coomassie blue and Silver staining.

### Deubiquitination assay

Deubiquitination assay was performed under denaturing conditions as described earlier with some modifications^18^. Briefly, cells were treated with the indicated amount of MG-132 for 4 hrs to allow accumulation of poly-Ub proteins before harvesting. Cells were then lysed in SDS-lysis buffer (50mm Tris-Cl pH 6.8, 1% SDS, 10% glycerol, 1mM Na_3_Vo_4,_ and protease inhibitor cocktail, calbiochem). After boiling for 10 min, lysates were diluted ten times with IP-Lysis buffer supplemented with 15mM NEM (Sigma) and protease inhibitor cocktail. The diluted lysates were immunoprecipitated with indicated antibodies and probed with anti-ubiquitin antibody. Finally, 3% input was run for the detection of other indicated proteins by IB.

### MTT assay

Cytotoxicity of USP7 inhibitor P5091 and P22077 was determined by MTT (3-(4,5-dimethylthiazol-2-yl)-2,5-diphenyl tetrazolium bromide) assay (Sigma-Aldrich, St. Louis, MO, USA). Cytotoxicity was assessed after 24 hrs and 48 hrs of inhibitor treatment in 96 well plates.

### Flow cytometry

For cell cycle analysis, the treated cells were harvested using Trypsin. The cells were ethanol (70% ice-cold EtOH) fixed and further processed for cell-cycle analysis as described before^17^. To investigate the magnitude of apoptosis induced by USP7 inhibition, cells at 70% confluency were treated with the indicated concentration of USP7 inhibitors for 24hrs and 48hrs. Apoptotic cell death was analyzed by staining with Annexin V/PI as per the manufacturer’s instructions (APOAF-20TST, Merck Millipore) and analyzed in BD LSR-Fortessa using FACS-Diva software (BD Biosciences). Floating dead cells were collected and added to the harvested cells for analysis.

### Caspase3/7 assay

HCT116 cells were plated in 96-well plates (in triplicate) and treated with DMSO or p5091 for 24 hrs. Caspase-Glo 3/7 Reagent (Caspase-Glo® 3/7 Assay, Promega) was added to the wells, and luminescence was recorded after 30 min, and also after 1.5 hrs, following manufacturer’s instructions.

### RNA preparation and quantitative real-time PCR (qRT-PCR)

Total RNA was extracted using Trizol reagent (Invitrogen) and processed for real-time PCR as previously described^19^. Briefly, cDNA was prepared, and Real-time PCR was performed using SYBR Green Master Mix (Bio-Rad) in Via7 Real-Time PCR Instrument (Applied Biosystems), described before. Sequences of the primers are given in Additional file 1.

### Immunofluorescence microscopy

Immunofluorescence procedure was described previously^20^. Briefly, HEK cells were seeded over sterile coverslips placed inside 35 mm tissue culture dishes and cultured as per recommended conditions to a confluency of 60%. The cells were processed as follows: 4% paraformaldehyde fixation, 0.5% Triton-X-100 permeabilization, and blocking with 2.5% BSA in PBS. A standard protocol for immuno-staining was followed, and the cells were stained with primary antibodies and fluorochrome-conjugated secondary antibodies (Alexa-Fluor 488 and/or 594). In addition, 4, 6-diamino-2-phenylindole (DAPI) was used as a nuclear counterstain. All slides were viewed, and images captured at 120X using FluoView FV10i confocal laser scanning microscope (Olympus Life Science).

### Colony formation assay

The cells (2.5×10^3^) in triplicate plates were treated with DMSO or USP7 inhibitor for 24 hrs. After treatment, the cells were grown for another 15 days in complete medium. The cells were then fixed with 4% paraformaldehyde and stained with freshly prepared 0.1% crystal violet stain for 10 min. Following rinsing with distilled water, the colonies that had formed in each well were counted and captured the images.

### Cell lysate preparation for proteomics analysis

Cell pellets from three replicates in each group were lysed mechanically with a needle in the absence of protease inhibitors in lysis buffer, as described before. Briefly, 0.3 mL of ice-cold PBS was added to frozen cell pellets, and the resulting mixture was lysed by passing through 23-gauge needles for 15 times, centrifuged at 15000g at 4^0^C to separate cytosolic protein fraction from debris. Protein concentration of the extracts was measured by Bradford assay (Bio-rad). A total of 20μg protein from each of the three replicates (control and treated) was digested with 1μg of sequencing grade trypsin (Promega, Fitchburg, WI) followed by DTT and iodoacetamide treatment. Following overnight digestion at 37°C, samples were acidified with 5% formic acid to stop Trypsin activity and prepared for LC−MS/MS by C18 Zip-Tip purification as per manufacturer’s protocol (Millipore, Billerica, MA). Peptide samples were resuspended in water with 0.1% formic acid (v/v) and analyzed by nano-LC−MS/MS.

### LC-MS/MS

For protein identification and label-free quantification, six replicates, three from each sample, were analyzed by nano-LC−MS/MS. For each run, 1μg of the digest was injected on a reverse-phase C18 column using Thermo Scientifics nano-LC system. Water (A) and acetonitrile (B), both with 0.1% formic acid used as chromatography solvents. Peptides were eluted from the column with the following gradient - 3 to 35% B (130 min). At 140 min, the gradient increased to 95% B and was held there for 10 min. At 160 min, the gradient returned to 3% to re-equilibrate the column for the next injection. A short 50 min linear gradient blank was run between samples to prevent sample carryover. Peptides were analyzed by data-dependent MS/MS on a Q-Exactive Orbitrap mass spectrometer (Thermo Fisher Scientific, MA). In brief, the instrument settings were as follows: the resolution was set to 70,000 for MS scans and 17,500 for the data-dependent MS/MS scans to increase speed. The MS AGC target was set to 10^6^ counts, while MS/MS AGC target was set to 10^5^. The MS scan range was from 300 to 2000 m/z. MS scans were recorded in profile mode, while the MS/MS was recorded in centroid mode to reduce data file size. Dynamic exclusion was set to a repeat count of 1 with a 25 s duration. Following LC−MS/MS acquisition, all the data were searched using Sequest HT Proteome Discoverer 1.4 search engine (Thermo Fisher Scientific) against the Uniprot Human database at a false discovery cut off ≤1%. After identification, the LC−MS/MS data were aligned using Chromalign. Quantitation of peptides in both samples was performed using SIEVE 2.1 (Thermo Fisher Scientific).

### Domain-wise docking of USP7 and XIAP

The 3D structures of TRAF (PDB id: 2F1W), catalytic domain (PDB id: 1NBF) and Ub like domain (PDB id: 2YLM) of USP7 protein, and BIR1 (PDB id: 2POI), BIR2 (PDB id: 1C9Q), BIR3 (PDB id: 1F9X), UBA (PDB id: 2KNA) and Ring (PDB id: 4IC3) domains of XIAP protein were extracted from protein databank (PDB)^21^. Each domain of USP7 was docked with every domain of XIAP using PatchDock^22^, clusPro^23^ and swarmDock^24^docking programs, respectively. Top 100 docking solutions for each pair were selected based on docking score and used for RMSD based clustering. The clustering was performed using UCSF chimera^25^, and largest cluster for each pair was selected and ranked based on the average docking score of the docking solutions present in largest cluster. Top 6 pairs based on rank were chosen from each of the docking programs and compared. The pairs commonly present in top six of all the three docking programs were selected as the probable interacting domains of USP7 and XIAP.

### Modelling and MD simulation of full-length USP7 and XIAP

The full-length structures of USP7 (amino acids: 61-1087) and XIAP (amino acids: 20-497) were modelled using Modeller v9.16^26^. The crystal structures of TRAF, catalytic and Ub like domains of USP7 (*PDB ids: 2F1W, 2F1Z, 5KYC, 2YLM and 4PYZ*), and BIR1, BIR2, BIR3, UBA, and RING domains (*PDB ids: 2POI, 1C9Q, 4J3Y, 1F9X, 2KNA and 4IC3*) with a *de novo* model (amino acids 98-125) predicted by robetta^27^ of XIAP were used as templates to model the full-length structures of USP7 and XIAP, respectively. The stereo-chemical properties and the fold compatibility of the final models (USP7 and XIAP) were validated by Rampage^28^, Prove^29^, Verify3D^30^, and Errat^31^ programs. The final models of USP7 and XIAP were subjected to molecular dynamic simulation to get the energetically stable structures. Molecular dynamic (MD) simulation of USP7 and XIAP proteins were carried out using Desmond^32^ simulation package. The OPLS_2005 force field parameters^33, 34^ were used for the simulation. The system was solvated with TIP3P^35^ water, and counter ions were added for neutralizing the system. Orthorhombic periodic boundary conditions were defined to specify the shape and size of the simulation box buffered at 10Å distances. After building the solvated system, minimization and relaxation of the system were performed under NPT ensemble using default protocol of Desmond, which consist of a total of 8 stages among which there are 2 minimization steps (restrained and unrestrained) and 4 short simulations (equilibration phase) steps before starting the actual production run. A hybrid method combining the steepest decent and the limited-memory Broyden-Fletcher-Goldfarb-Shanno (LBFGS) algorithm was used for minimizing the system. The temperature of NPT-simulations was regulated with Nose-Hoover Chain thermostat^36^ with 1.0 picoseconds (ps) relaxation-time, and the pressure was controlled with Martyna-Tobias-Klein barostat^37^ with isotopic coupling and 2.0 ps relaxation time. The final production run was carried out under normal pressure and temperature (NPT) conditions for 300 ns at 300K temperature and 1 atmospheric pressure with a time step of 2 fs. The RESPA^38^ integrator was used for integrating the equations of motion. The trajectories were saved at the interval of 4.8 ps. Simulation Quality Analysis (SQA) program was used to calculate the Energies of the protein during the simulation time. Root Mean Square Deviation (RMSD) and Root Mean Square Fluctuation (RMSF) were calculated using simulation event analysis (SEA) programs available with the Desmond package.

### Molecular docking and MD simulation of the docked complex

The stable USP7 and XIAP structures of full length USP7 and XIAP proteins obtained after simulation were docked using the protein-protein docking mode of SwarmDock^24^ docking program. The probable docking pose was selected based on the docking score. The selected docked USP7-XIAP complex was further simulated for 100 ns to check the stability of the complex. The USP7-XIAP complexes (before and after simulation) were validated by estimating the true-like complex probability at PCPIP (Protein Complex Prediction by Interface Properties) server^38^ and MM/GBSA based binding energy estimation by MM/GBSA module of HawkDock server^39, 40^.

### Preparation of nanoencapsulated USP7 inhibitors P5091 and P22077

A modified emulsion-diffusion-evaporation method was used to formulate the nanotized P5091 and P22077 as described before with some minute modification in drug and polymer ratio. In brief, 50mg of PLGA (50:50, MW: 7,000–17,000) was dissolved in 500 ul of ethyl acetate, and 5mg of the commercially available pure P5091 or P22077 were added to this mixture. The organic solution was then emulsified with an aqueous phase containing di-dodecyl-di-methyl ammonium bromide (DMAB). The resulting oil-in-water emulsion was stirred at room temperature for 3 hr before homogenizing at 15,000 rpm for 5 minutes. Subsequently, the organic solvent was removed by rotary evaporation and the aqueous phase containing nanoparticles was sonicated for 30 minutes. The resulting aqueous emulsion was centrifuged at 35,000 rpm for 1 hr and the nanoprecipitate was dried and suspended in phosphate buffered saline (PBS), along with 20% sucrose and glucose (used as a cryoprotectant), lyophilized and stored at room temperature for future use.

### Physical characterizations of nanoformulations

#### Particle size measurement

the particle size of nanotized USP7 inhibitors P5091 and P22077 as well as empty nanoparticles were measured by Nanosizer 90 ZS (Malvern Instruments, Southborough, MA). The samples were prepared by taking 1mg of each nanoformulations in 10 ml of ddH_2_O and the particle size were accurately measured by dynamic light scattering (DLS). Measured particle size was represented as average of 10 runs with triplicate measurement in each run. Zeta potential of the synthesized nanoparticles was measured by Zetasizer Nano ZS (Malvern Instruments, Southborough, MA, USA). Prior to analysis, the solution in ddH_2_O was filtered through a 2μm filter. Each sample was measured in triplicate.

#### Atomic Force Microscopy (AFM)

The size and shape were measured by atomic force microscopy using Agilent Technologies 5500 Pico Plus AFM system. All the images were obtained in the aquatic mode using cantilevers having a resonance frequency of 146–236 kHz, tip height 10–15 μm, and tip length 225 μm. Mica was chosen as a solid substrate and used immediately after cleavage in a clean atmosphere. During the characterization experiment, the probe and cantilever were immersed completely in the water solution. The nanocapsule suspensions on mica were dried in air (65% humidity) for 30 min. Images were captured and analyzed using Picoscan 5.33 software from Molecular Imaging Corp.

#### Transmission Electron Microscopy (TEM)

The size and morphology of P5091 and P22077 nanoformulations were examined using a JEOL JEM-2100 transmission electron microscope (JEOL, Inc., Peabody, MA, USA) at an acceleration voltage of 300 kV. One drop of nanoparticle suspension was dispersed in deionized water, lyophilized and mounted on a thin film of amorphous carbon deposited on a copper grid (300 meshes). After drying under clean condition, the grid was examined directly using TEM.

#### Entrapment efficiency and in vitro release study

Centrifugation method was performed to determine the encapsulation efficiency (EE). The particles were precipitated. A certain proportion of fresh nanoformulations was dissolved in ethyl acetate. The entrapment efficiency was measured by using a UV-VIS spectrometer. The drug was detected at 320 nm. The EE parameter was calculated as follows: EE = Wt – Wd/Wt. Where Wt and Wd describes the total DIM added and the extracted molecule into the supernatant, respectively.

*In vitro* release profiles of P5091 and P22077 from PLGA nanoparticles were determined in phosphate buffer saline (PBS; pH 7.4) where release occurred till 80 hrs.

#### FTIR study

The effects of encapsulation process on the chemical group and the interaction between the components was studied by performing a Fourier Transform Infrared Spectroscopy (FTIR) model Spectrum TWO (Perkin Elmer, USA). The FTIR spectra ranging between 500 cm^-1^ and 4000 cm^-1^ was obtained by mixing samples with KBr powder (Infrared grade).

#### TG-DTA analysis

The thermal stability of the nanoformulations were confirmed by simultaneous TG and DTA. The curves were obtained in the temperature range from 25 to 500°C, using aluminium crucibles with about 5 mg of samples, under dynamic air atmosphere (100mL/min) and heating rate of 20°C/min.

### Statistical analysis

Statistical analysis of results was performed by Student’s t-test, or one-way ANOVA, as specified in the individual figure legends according to the assumptions of the test using GraphPad Prism software 8.4.2. The bars shown represent the mean ± standard deviation (SD). The P-values are presented in figure legends where a statistically significant difference was found. *P < 0.05; **P < 0.01; ***P < 0.001, ****P<0.0001.

## Results

### Induction of p53 independent apoptosis upon USP7 inhibition: XIAP is a possible mediator

It is well established that apoptosis induction due to pharmacological inhibition of USP7 is mainly through the MDM2-p53 axis^41^. However, several other reports indicate that p53 independent apoptosis can also occur due to induction of oxidative and endoplasmic reticulum stress upon inhibition of USP7^42^. Consistent with this observation, we found no significant difference in cell viability upon P5091 (a potent USP7 inhibitor) treatment in HCT116 cells having either p53^Wt^ or p53^Null^ status, as there was no significant difference in IC-50 values at 24 and 48 hrs (Figure 1A). A similar result was also found when we checked the degree of apoptosis by FACS analysis (Supplementary Figure 3A). Additionally, cleavage of PARP and Caspase3 in P5091 treated p53^Wt^ and p53^Null^ HCT116 cells indicated that the apoptosis was independent of cellular p53 status (Figure 1B). P5091 treatment in a dose-dependent manner resulted in a significant increase in active caspases in the HCT116 p53^Null^ cell line (Figure 1C). Additionally, reduction in colony formation ability upon USP7 inhibition was also independent of cellular p53 status, as similar results were observed in HCT116 (p53^Wt^), HepG2 (p53^Wt^), and in Huh7 (p53^Mut^) cell lines (Figure 1D). Thus, all these results indicate that magnitude of apoptosis induction due to USP7 inhibition in p53^Null^ HCT116 cells was similar to the p53^Wt^ HCT116 cells. Hence, we concluded that there was some unidentified pathway at play that inducing apoptosis upon USP7 inhibition which is independent of cellular p53 status. To investigate this hypothesis, we used an affinity-based purification method followed by LC-MS/MS. Purified recombinant GST-USP7 was used as bait to pull USP7 interactome from HEK whole cell lysate (WCL). Eluted proteins were separated by SDS-PAGE, gel was silver-stained (Figure 1E), and further processed and finally analyzed by mass spectrometry using nano LC-MS/MS to identify the USP7 interacting proteins. Empty bead and GST were used as controls. A list of identified interacting proteins was given in supplementary Table 1. To further identify the bonafide USP7 interactors from the initial screen, total spectral counts from GST-USP7 pulldown proteins were compared with bead and GST control as well as to the contaminant repository for affinity purification (*CRAPome* http://www.crapome.org/)^43^ and selected based on SAINT score^44^. The topmost USP7 interacting proteins were identified from two independent experiments, are shown in Table 1. We were able to locate multiple USP7 target proteins. Among them, XIAP presented itself as a promising candidate with a significant SAINT score. USP7 and XIAP interaction was again verified by a reverse pulldown assay where we used recombinant GST-XIAP in affinity chromatography to find out XIAP interactome from HEK WCL (Figure 1F). A detailed list of XIAP interacting proteins obtained from this study was given in Table 2 and Supplementary Table 2.

**Figure 1.**
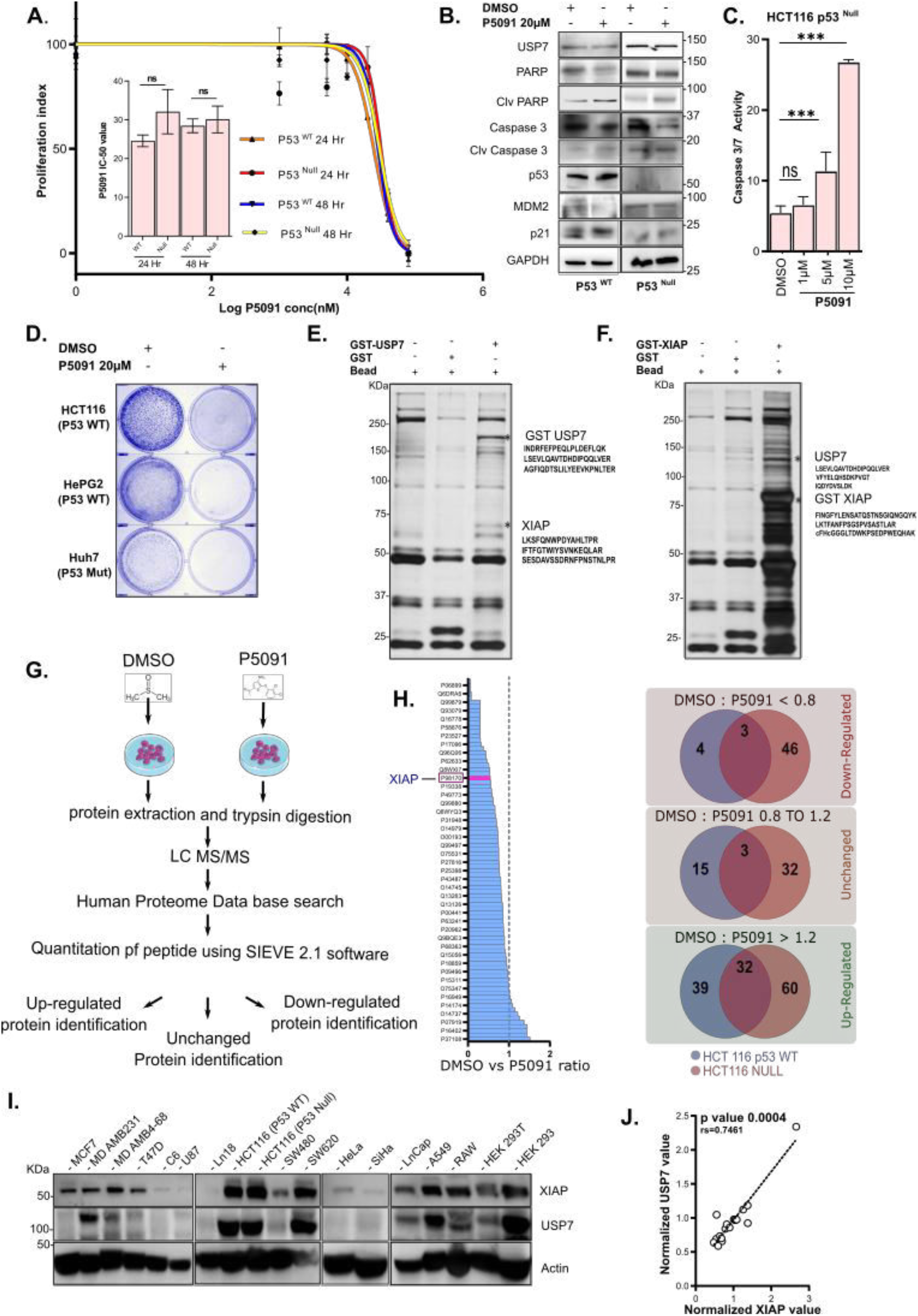
P53 independent apoptosis by USP7 inhibition: XIAP is a probable candidate. **A,** HCT116 (p53^Wt^ and p53^Null^) lines were treated with the indicated dose of P5091 (0 to 80 µM) for 24 and 48 hrs, and cell viability was checked by MTT assay. Inset graph representing change in mean IC-50 value of both cell lines at different time points. **B,** Protein levels of USP7, PARP, Cleaved PARP, Caspase3, Cleaved Caspase3, p53, MDM2 and p21 were determined by immunoblotting (IB) where HCT116 (p53^Wt^ and p53^Null^) cell lines were treated with USP7 inhibitor P5091 (20µM) for 24 hrs. GAPDH was kept as loading control. **C,** Following treatment of HCT116 (p53^Null^) cells with P5091 in different doses (1, 5, 10µM) for 24 hrs, the increase in caspase 3/7 activity was determined using a fluorescence plate reader. Data represent mean ± SD of three independent biological replicates. **D,** Number of colonies formed by HCT116 (p53^Wt^), HepG2 (p53^Null^), and Huh7 (p53^Mut^) cells after treatment with P5091 (20 µM) for 24 hrs; colonies were counted after 15 days. **E & F,** Identification of USP7 and XIAP interacting proteins by Mass Spectrometry. Silver stained SDS-PAGE gels containing elute from respective pull-down as indicated. MS analysis identifies specific peptides of XIAP and USP7 respectively from GST-USP7 and GST-XIAP pull-down lanes by using HEK cell lysates. **G,** Identification of USP7 target proteome in HCT116 cells by label-free comparative proteomics analysis upon USP7 inhibition with P5091 (15µM for 24 hrs), figure represents the workflow of label-free quantitation (LTQ) by nano-LC−MS/M.S. on a Q-exactive followed by SIEVE^TM^ processing. **H,** Graphical representation of calculated protein ratios of P5091 treated samples with respect to the vehicle control (left panel). Venn diagram showing identified number of Up-regulated, Unchanged and Down regulated proteins in HCT116 p53^wt^ and HCT116 p53^null^ cell lines upon P5091 treatment with respect to control (right panel). **I & J,** Whole-cell lysates were prepared from MCF7, MD-AMB 231, MD-AMB 468, T47D, C6, U87, LN18, HCT116 (p53^Wt^), HCT116 (p53^Null^), SW480, SW620, HeLa, SiHa, LnCap, A549, RAW, HEK293T, and HEK293 cells. IB analysis was performed using respective antibodies for USP7, XIAP and β-Actin. Normalized values were plotted to show a strong positive correlation between USP7 and XIAP where p value < 0.0001, n=2. Error bars in all the indicated sub-figures represent mean ± SD from three independent biological repeats. Indicated P-values were calculated using Student’s t-test and P<0.05 is represented as *, otherwise non-significant (ns).

**Table 1.** USP7 interactome in HEK293T cells

| Uniprot Ac. No. | Gene Name | FC-A value | SAINT Score |
| --- | --- | --- | --- |
| Q93009 | **USP7** | 125.16 | 1 |
| P78347 | GTF2I | 74.16 | 1 |
| P09874 | PARP1 | 74.16 | 1 |
| P52272 | HNRNPM | 54.21 | 1 |
| P98170 | **XIAP** | 49.78 | 1 |
| P16152 | CBR1 | 40.91 | 1 |
| Q86U86 | PBRM1 | 40.91 | 1 |
| O75643 | SNRNP200 | 34.26 | 1 |
| Q7L2E3 | DHX30 | 32.04 | 1 |
| O00571 | DDX3X | 29.6 | 1 |
| Q9NZI8 | IGF2BP1 | 28.86 | 1 |
| Q8IX18 | DHX40 | 25.39 | 1 |
| P43243 | MATR3 | 25.39 | 1 |
| Q16643 | DBN1 | 23.17 | 1 |
| P42166 | TMPO | 23.17 | 1 |
| Q92922 | SMARCC1 | 23.17 | 1 |
| Q02218 | OGDH | 23.17 | 1 |
| Q6P2Q9 | PRPF8 | 23.17 | 1 |
| Q9BQG0 | MYBBP1A | 20.95 | 1 |
| Q15029 | EFTUD2 | 20.95 | 1 |

**Table 2.**
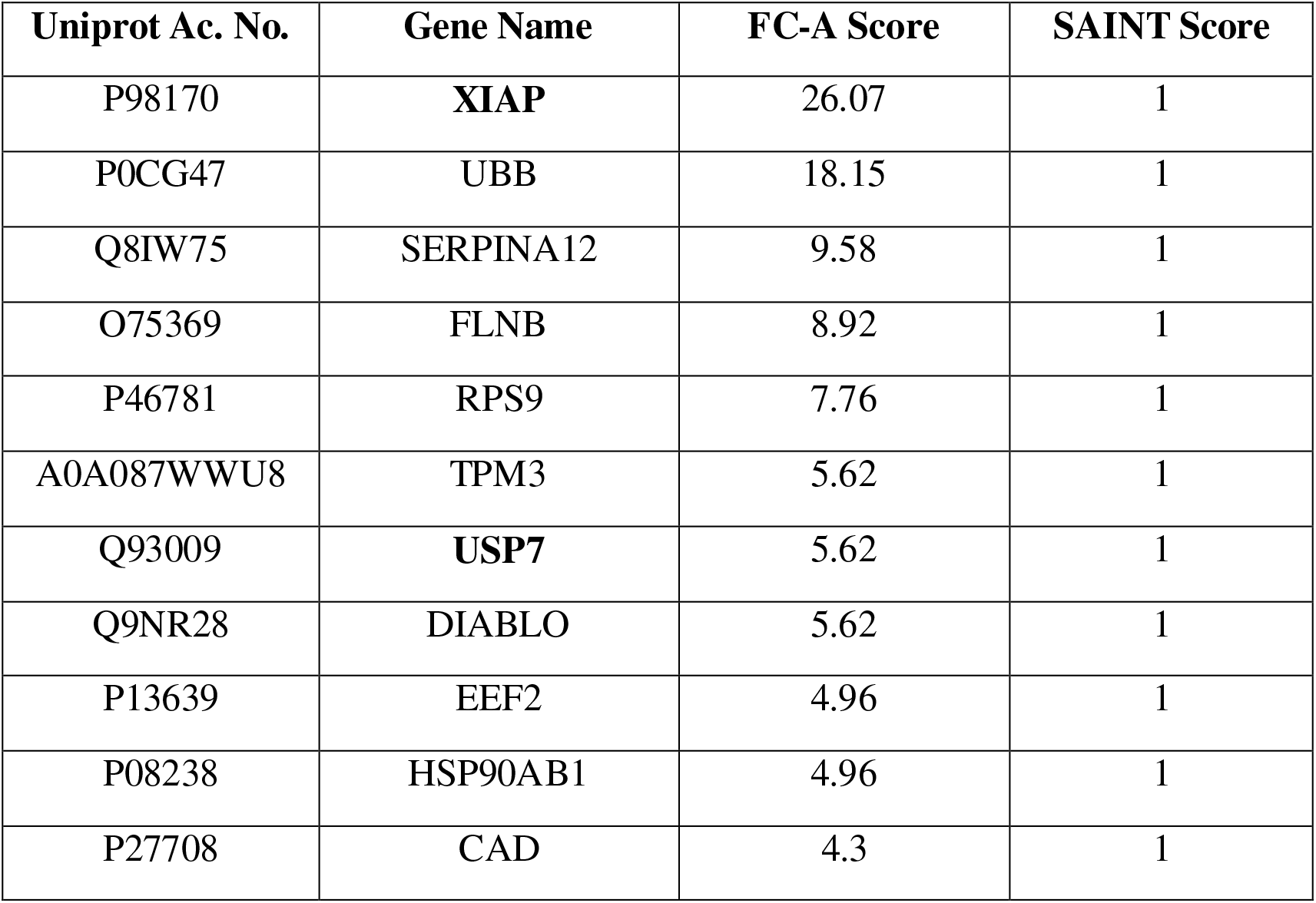
XIAP interactome in HEK293T cells

| **Uniprot Ac. No.** | **Gene Name** | **FC-A Score** | **SAINT Score** |
| --- | --- | --- | --- |
| P98170 | **XIAP** | 26.07 | 1 |
| P0CG47 | UBB | 18.15 | 1 |
| Q8IW75 | SERPINA12 | 9.58 | 1 |
| O75369 | FLNB | 8.92 | 1 |
| P46781 | RPS9 | 7.76 | 1 |
| A0A087WWU8 | TPM3 | 5.62 | 1 |
| Q93009 | **USP7** | 5.62 | 1 |
| Q9NR28 | DIABLO | 5.62 | 1 |
| P13639 | EEF2 | 4.96 | 1 |
| P08238 | HSP90AB1 | 4.96 | 1 |
| P27708 | CAD | 4.3 | 1 |

Next, in an attempt to identify the targets of USP7, we performed a label-free quantitation (LFQ) of USP7 inhibited HCT116 cell lysates. For this, HCT116 (p53^WT^) cells were treated with P5091 (15µM) for 24 hrs. Data acquired by nano LC-ESI-MS/MS on an Orbitrap LTQ spectrometer were processed with SIEVE^TM^ 2.1 software to reveal up-or down-regulated proteins^45^ (Figure 1G). To maximize the statistical relevance of our results, we analyzed the data generated from three independent biological replicates. Following LC-MS/MS, SIEVE alignment of the replicates MS intensity against retention time was carried out using the proprietary algorithm Chromalign. We successfully identified 34 upregulated and 46 down-regulated proteins (Figure 1H, Supplementary Table 3). Identified proteins with significant fold changes were further analyzed for KEGG and Reactome Pathway enrichment and also for GO Biological Process and GO Molecular Function (Supplementary Figure 1 and Supplementary Figure 2) in Cytoscape Visualization tool (see Material & Methods). The results from our proteomic analysis revealed XIAP as a potential interacting partner of USP7, and it is downregulated with a *p* value of 0.0246 upon USP7 inhibition. So, it can be considered as a significant target of USP7.

Furthermore, expressions of XIAP and USP7 proteins were checked in multiple cell lines of breast, glioblastoma, colorectal, cervical, prostate, lung cancers, macrophage, RAW, HEK293, and HEK293T cells. A strong positive correlation between XIAP and USP7 expressions was observed (*p* values < 0.00001) (Figure 1I & J). To further strengthen our results, we analyzed the expression pattern of USP7 and XIAP from data obtained from ProteomicsDB^46^ (www.proteomicsdb.org). We also found a positive correlation between USP7 and XIAP expressions, where r_s_= 0.21, p=0.02, and n= 120 (Supplementary Figure 3B). We also checked for expression correlation in cells originated from different sources viz., Skin, Lung, Colon, and Breast tissues show a significant positive correlation, on the contrary cells originated from the ovary, brain, circulatory system show a negative correlation of their expressions pattern (Supplementary Fig 3C). List of cells analyzed for this purpose given in Supplementary information (Supplementary Table 4).

### Identification of XIAP as a putative substrate of deubiquitinase USP7

Deubiquitinating enzymes can efficiently regulate the stability of target proteins by removing ubiquitin (Ub) moieties from their ubiquitinated substrates. To begin assessing the role of USP7 deubiquitinase activity on XIAP stability, we transiently overexpressed GFP-USP7 in HEK cells (Figure 2A Lanes 1&2) and checked XIAP protein levels. A significant increase in XIAP level was observed due to USP7 overexpression. On the other hand, the catalytic mutant of USP7 (C223S: designated as USP7^C223S^) decreased XIAP levels (Figure 2a lanes 3&4). Concomitantly, upon USP7 knockdown with three independent shRNAs of USP7 in HEK cells, a reduction in XIAP protein level was observed (Figure 2A lanes 5 to 8). USP7 knockdown using small interfering RNA (siRNA) resulted in a similar decrease in XIAP protein level; however, XIAP transcript levels remain unchanged. Similarly, when we treated HEK cells with the potent USP7 inhibitor – P5091^47^, a significant decrease in XIAP protein level was observed, but no detectable changes in XIAP transcript levels (Figure 2B). Thus, USP7 overexpression did not have any role in changing *XIAP* gene transcription.

**Figure 2.**
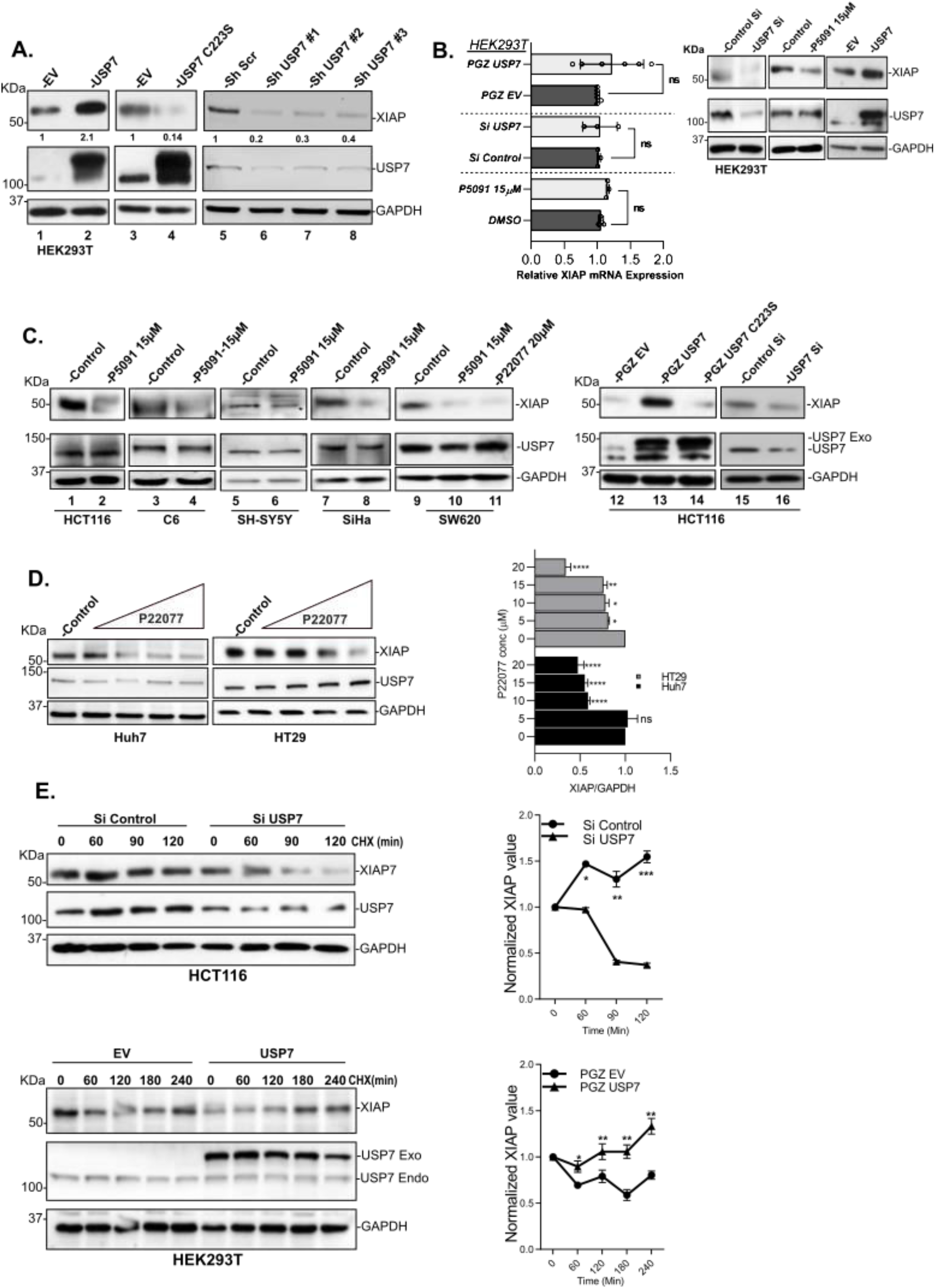
XIAP is a putative substrate of USP7. **A,** HEK cells transfected with 2µG of EV (pGZ) (lane-1&3), pGZ-USP7 (lane-2), and consecutively inactive form of USP7 mutant pGZ-USP7^C223S^ (lane-4); 48 hrs post-transfection, USP7 and XIAP protein levels were checked by IB. Additionally, three different shRNAs targeting USP7 and scramble shRNA were transfected (lanes-5,6,7&8) and the levels of XIAP and USP7 were analyzed by IB. GAPDH was kept as loading control. **B,** qRT-PCR analysis showing relative mRNA fold change upon USP7 inhibition by P5091 (15µM) for 24 hrs. Relative mRNA fold change of XIAP upon USP7 knockdown (using siUSP7) and USP7 over-expression. Protein expression of XIAP from the same experiment was also represented by IB, where GAPDH was used as an internal loading control. **C,** Multiple cancer cell lines such as HCT116, C6, SH-SY5Y, SiHa, and SW620 were treated with 15µM of USP7 inhibitor (P5091-lane 2, 4, 6, 8, 10; P22077-lane 11) for 24 hrs and XIAP was analyzed by IB and compared with respective control lanes (lane 1, 3, 5, 7, 9). IB analysis of XIAP protein level in HCT116 cells transfected with GFP-USP7 (4 µG, lane 13), pGZ-USP7^C223S^ (4 µG, lane 14), or siUSP7 (lane 16). EV (lane 12), control Si (lane 15) were kept for control. **D,** Huh7 and HT29 cells were treated with USP7 inhibitor P22077 in a dose-dependent manner (5, 10, 15, 20 µM) for 24 hrs. XIAP was checked by IB. GAPDH was kept as an internal loading control. **E,** HCT116 cells were transfected with either Control Si (upper left panel) or siUSP7 (Upper right panel) and HEK cells were transfected with either EV (Lower left panel) or GFP-USP7 (Lower right panel) before treatment with cycloheximide (CHX: 50µG/ml). XIAP protein levels at indicated time points were analyzed by IB as shown in the figure (left panels). Graph showing the change in XIAP half-life was determined in USP7 knockdown and overexpression conditions. Error bars in all the indicated sub-figures represent mean (±) SD from three independent biological repeats. Indicated P-values were calculated using Student’s t-test and P<0.05 is represented as *, otherwise non-significant (ns).

Next, we sought to check the universality of XIAP regulation by USP7. For this, we examined this phenomenon in different cancer cell lines. We used two well-established small molecule inhibitors of USP7 *viz*., P5091 to inhibit USP7 in HCT116 (colorectal cancer), C6 (rat glioma), SH-SY5Y (neuroblastoma), and SiHa (cervical cancer) cells and checked XIAP protein levels (Figure 2C, lanes 1-8). Furthermore, USP7 was inhibited by both p5091 and P22077^48^ in SW620 (colorectal cancer) cells to see the expression of XIAP (Figure 2C, lanes 9-11). In all the cancer cell lines, USP7 inactivation led to a significant decrease in XIAP protein level. HCT116 cells transiently transfected with GFP-USP7 and its catalytic inactive mutant (USP7^C223S^) exhibited an increase in XIAP protein level upon overexpression of wild type USP7, but the opposite effect was observed in the case of mutant USP7 and in knocked down of USP7 by siUSP7 (Figure 2C, lanes 12-16). Additionally, USP7 inhibition by P22077 reduced XIAP protein levels in a dose-dependent manner in both Huh7 and HT29 cells (Figure 2D). all these data strongly suggest that XIAP is regulated post-transnationally by USP7 in several cancers. Furthermore, the half-life of endogenous XIAP was monitored and found to be decreased significantly upon USP7 knockdown in HCT116 cells (Figure 2E Upper panel). The reverse was observed in case of USP7 overexpression (Fig 2E lower panel). These results indicate that the change in the XIAP protein level may be due to its stabilization by the USP7.

### USP7 and XIAP colocalize and physically interact *in vitro*

To justify our observation, next we investigated the molecular mechanism underlying the stabilization of XIAP by USP7. Being a deubiquitinase, we speculated the upregulation of XIAP at protein level by USP7 was mainly due to its deubiquitinase activity, and hence the physical interaction between the two proteins is necessary. Initially, we checked the subcellular localization status of both the proteins by immunocytochemistry (ICC) in HEK cells. USP7 showed a multi-compartment localization within the cell. So far, USP7 has been demonstrated to be localized primarily in the nucleus and associated with PML nuclear bodies and ND-10^49^. Additionally, the presence of USP7 in the cytoplasm^50, 51^ and in mitochondria^52^ has been reported. In our study, confocal imaging of HEK cells showed abundant yellow spots (USP7-XIAP co-localization) mainly in the cytoplasm, revealing that both proteins showed cytoplasmic co-localization *(Pearson’s correlation Coefficient, r=0.604)* (Figure 3A). Next, we attempted to establish the nature of USP7 and XIAP interaction. For this, we purified and incubated GST-XIAP with Myc-His-USP7 *in vitro*. Pull-down experiments were performed with glutathione conjugated protein A sepharose beads, and eluted proteins were separated by SDS-PAGE. The presence of proteins was analyzed by Coomassie blue staining. The result revealed the USP7 band at the GST-XIAP lane but not in the control GST lane indicating a direct interaction between USP7 and XIAP in the purified system (Figure 3B). Similarly, recombinant GST-XIAP can pull endogenous USP7 from HEK lysate where we kept GST as control (Figure 3C).

**Figure 3.**
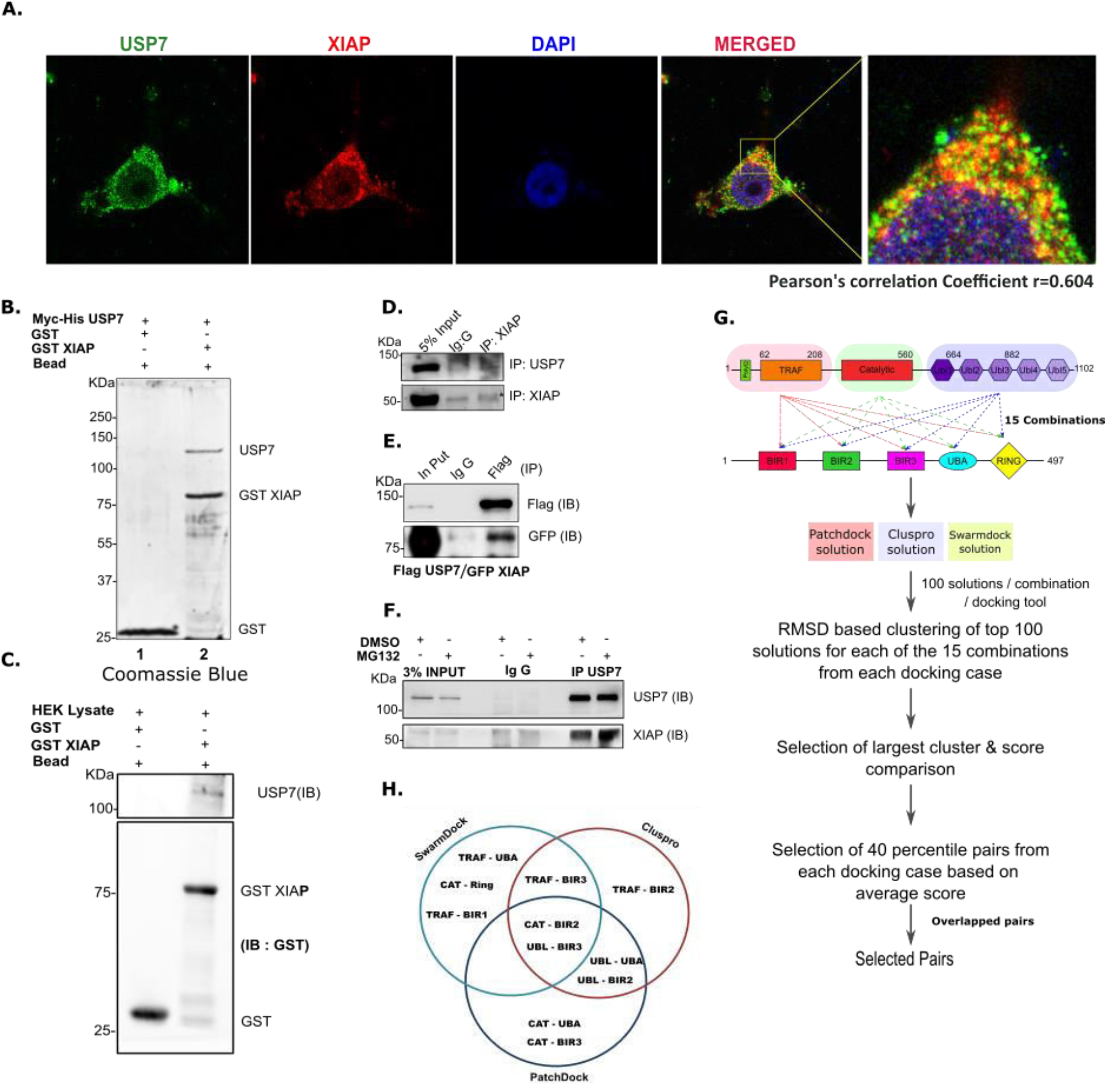
USP7 and XIAP colocalized and physically interacted in *in vitro* purified system and *ex-vivo* in cell lysates - identification of their molecular motifs responsible for their interaction. **A**, Confocal images of HEK cells showing stains for USP7 (green), XIAP (red), and USP7-XIAP merged (yellow). Cells were counterstained with DAPI (blue). Images were captured at 60X optical magnification. **B,** Purified Myc-His USP7 incubated separately with purified GST and GST-XIAP in the presence of GST-Bead for 4 hrs. Proteins were separated by SDS-PAGE and analyzed by Coomassie blue staining to visualize the specific bands for GST (lane 1), GST-XIAP and Myc-His USP7 (lane 2). Figure shows the representative data of three biological replicates. **C,** HEK cell lysate was incubated separately with purified GST and GST-XIAP in the presence of GST-bead for 4 hrs, followed by IB using anti-USP7 and GST antibody. **D,** HEK cell lysate was immunoprecipitated with XIAP antibody followed by IB using antibodies against USP7 and XIAP to identify their interaction at the endogenous condition. **E,** HEK cells were transiently transfected with FLAG-USP7 and GFP-XIAP. Equal amounts of cell lysates were used for immunoprecipitation with Flag antibody and Normal Rabbit Serum, followed by IB using antibodies against Flag and GFP to identify their interaction under overexpressed conditions. **F,** HEK cells were treated with either DMSO or MG132 (25µM) for 8 hrs. Equal amounts of lysates were used for IP using USP7 antibody followed by IB using indicated antibodies to show an increased interaction of USP7 and XIAP upon blocking the proteasomal system. Veriblot secondary antibody was used to prevent nonspecific detection of the heavy and light chains. **G,** Workflow of domain wise docking of USP7 and XIAP proteins using three independent Docking software to identify the probable domains responsible for interaction. **H,** Venn diagram showing domain pair of USP7 and XIAP by using three docking softwares and identified CAT-BIR2 and UBL-BIR3 as the probable interacting domains.

Furthermore, in a reversed pulldown experiment using antibody against XIAP showed co-immunoprecipitation of endogenous USP7 (Figure 3D). We also confirmed by a reverse immunoprecipitation assay in a dual overexpression systerm (Figure 3E). Next, we increased the poly-ubiquitinated protein pool in HEK cells by treatment with MG132, and there was a significant increase in XIAP pulldown was observed (Figure 3F). All these results indicate that USP7 physically interacts with XIAP and this interaction is influenced by ubiquitination status of XIAP.

Next, to further decode the nature of USP7 and XIAP interaction in a domain-wise manner, we opted for computational modelling of protein-protein interactions through docking experiment as represented in a schematic diagram (Figure 3G). The crystal structures of each domain of USP7 (TRAF, catalytic, and Ub like domains) and XIAP (BIR1, BIR2, BIR3, UBA, and Ring-domains) were available in the protein databank (PDB). However, full-length structures were not available. Therefore, initially domain-wise docking of USP7 and XIAP was performed by three different standard docking programs followed by RMSD based clustering. The USP7 and XIAP domain pairs from each program were ranked based on the average docking score of the solutions in the largest cluster. The top six USP7-XIAP pairs were UBL-UBA, CAT-UBA, UBL-BIR2, CAT-BIR2, UBL-BIR3, and CAT-BIR3 when docking was performed using PatchDock. TRAF-BIR3, UBL-BIR2, TRAF-BIR2, CAT-BIR2, UBL-BIR3, and UBL-UBA were the top six USP7-XIAP pairs when docking was performed using ClusPro docking program. However, SwarmDock resulted in TRAF-UBA, CAT-BIR2, TRAF-BIR3, CAT-Ring, TRAF-BIR1, and UBL-BIR3 as top six USP7-XIAP pairs (Supplementary Figure 4A). CAT-BIR2 and UBL-BIR3 were common when the top six USP7-XIAP pairs were compared and considered the probable interacting domains of USP7 and XIAP (Figure 3H & Supplementary Figure 4B & 4C).

### Identification of molecular motifs of USP7 and XIAP responsible for interaction

To get a clear picture docking experiment was immediately performed using full-length structures of USP7 and XIAP modelled in-silico (see methods) as represented in the schematic diagram (Figure 4A). The probable binding pose was selected based on the docking score (Figure 4B). Then the selected USP7-XIAP complex was subjected to MD simulation for 100 ns. Favourable structural adaptation in the USP7-XIAP complex was observed. The probability of complex formation was found to be increased from 77 % (before simulation) to 84 % (after simulation), and the MM/GBSA binding energy was decreased from −52.39 Kcal/mol (before simulation) to −136.59 Kcal/mol (after simulation) suggesting better stability of the complex after simulation. RMSD plot suggests that the overall structure does not deviate too much (<2 Å) in the last 70-75 ns, and the overall energy indicates the system remains unchanged. There were least fluctuations in the interfacing residues suggesting the stability of the complex during simulation (Supple Figure 5A & 5B). The catalytic (CAT) and Ub like domains 123 (UBL123) of USP7, and BIR1, BIR1/BIR2 inter-region, BIR2, BIR2/BIR3 inter-region, BIR3, and UBA domains of XIAP were found to exist at the interface. The interface analysis suggested that the catalytic domain of USP7 was largely in contact with the inter-region of BIR1 and BIR2 of the XIAP protein. However, the UBL123 domain of USP7 was in contact with BIR2, inter BIR2/BIR3 region, BIR3, and few residues of UBA domains of XIAP (Figure 4B & Supplementary Figure 5C & D).

**Figure 4:**
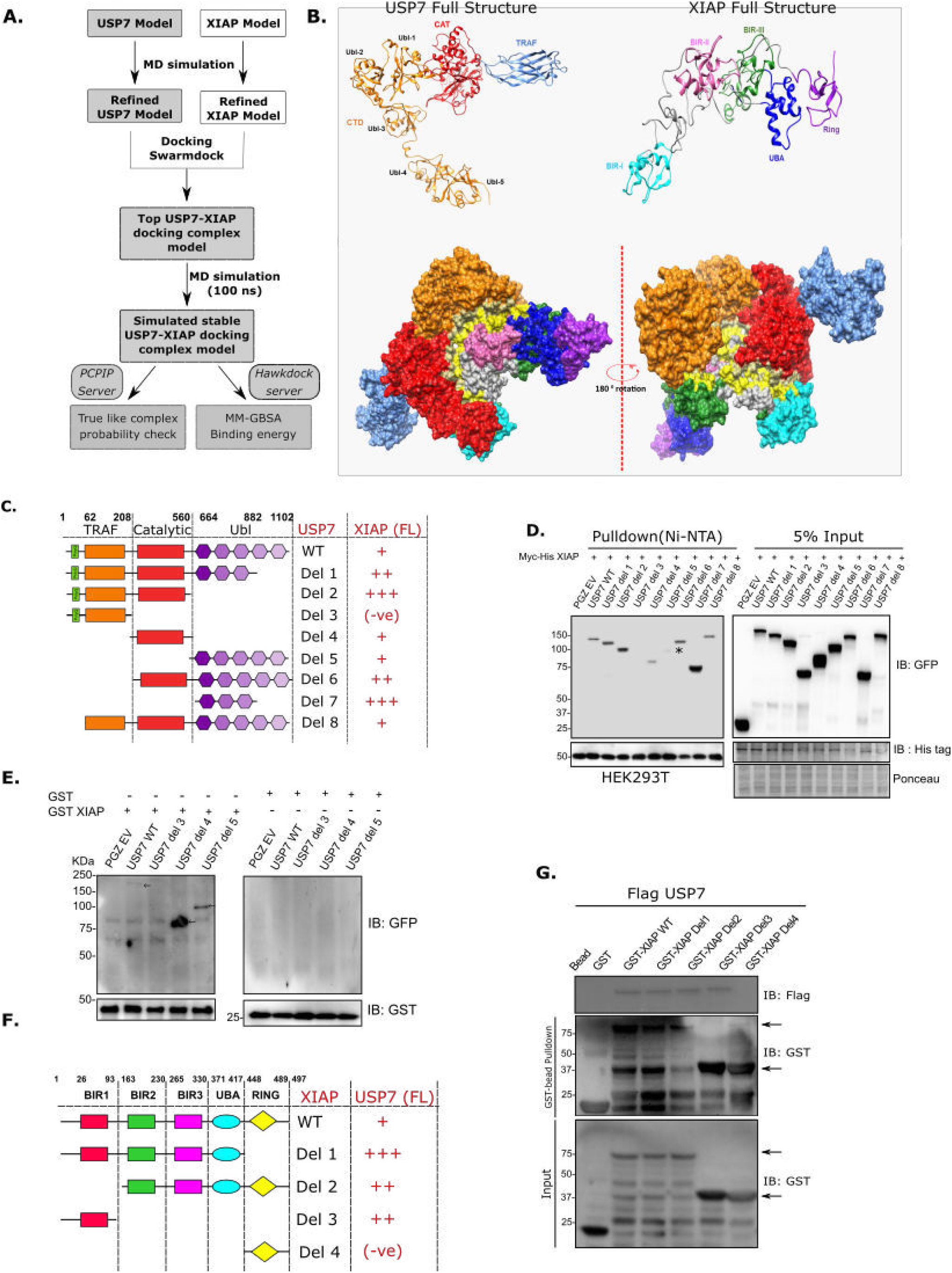
Bioinformatics analysis-based simulation of USP7 and XIAP and validation of identified molecular motifs of USP7 and XIAP responsible for their physical interaction. **A,** Workflow representing generation of full-length USP7 and XIAP followed by docking with Swarmdock and simulation of docking complex to identify the stable orientation of USP7 and XIAP interaction. **B,** Upper panels showing full-length USP7 and XIAP proteins generated by stitching the available USP7 and XIAP domains in PDB. Lower panels showing USP7-XIAP complex, where interface region highlighted in yellow. **C,** Schematic representation of USP7 deletion mutants and their interactions with full-length XIAP protein. **D,** HEK cells were co-transfected with indicated plasmid constructs; prepared the cell lysates followed by pull-down with Ni-NTA beads for 1 hr at room temperature. The pulled-down proteins and input were analyzed by IB with the indicated antibodies. Ponceau S staining indicates an equal loading in input lanes. HEK cells were transfected with the indicated plasmid constructs before the preparation of cell lysates. **E,** Equal amounts of lysate from plates transiently transfected as indicated, were incubated separately with purified proteins GST (right panel) or GST-XIAP fusion protein (left panel) including glutathione beads followed by IB analysis using indicated antibodies. The experiment performed thrice for biological replicates. **F,** Schematic representation of XIAP deletion mutants and their interaction with full-length USP7 protein. **G,** HEK cells were transiently transfected with Flag-USP7. The lysate was prepared and incubated separately with purified different GST tagged XIAP deletion mutants and GST protein (Control) overnight at 4°C. After pull-down with GST-bead for 2 hrs at RT, pulled-down proteins were analyzed by IB.

To validate our results generated from *in silico* analysis, we further characterized their interaction in HEK cells *in vitro*. For this, we developed specific deletion mutants of USP7, as depicted in the figure (Figure 4C). To identify the specific domains of USP7 responsible for its interaction with a specific region of XIAP, we conducted a Ni-NTA pulldown assay using cell lysates of HEK transfected with His-XIAP and individual USP7 deletion mutants as depicted in the figure along with full-length USP7 keeping EV as control. From this pulldown assay, we could conclude that the USP7 Ubl domain was mainly responsible for interaction with XIAP, where the presence of catalytic domain somehow influences their interaction, which concurs with our *in silico* data (Figure 4D). This was further fortified by GST-XIAP pulldown with USP7 deletion mutants, where mutants carrying only catalytic and Ubl domains showed positive interactions (Figure 4E). In a reverse approach, a similar pulldown experiment was performed with XIAP deletion mutants (based on our *in-silico* results), where we found BIR domains but not RING domain were responsible for interaction with USP7 (Figure 4F). Here, purified GST-tagged deletion mutants of XIAP were used to pull down FLAG-tagged USP7 overexpressed HEK cell lysate, where we found XIAP deletion mutants containing BIR and UBA domains able to interact with USP7 but XIAP mutants containing only RING domain unable to pull down USP7 (Figure 4G).

### USP7 protects XIAP from ubiquitination-dependent degradation in p53 independent manner

The next question, which we explored, was whether the interaction between USP7 and XIAP leads to the stabilization of XIAP. Blocking of proteasomal pathway mediated degradation of cellular proteins by MG132 treatment leads to a significant increase in the Poly-Ub-XIAP pool compared to the control, indicating the proteasome-mediated degradation of XIAP caused either by auto-ubiquitination or by involvement of other E3 ligases (Figure 5A). To understand the role of USP7 in the reversal of XIAP-ubiquitination, we performed an i*n vitro* deubiquitination assay. When USP7 was inhibited by P5091 treatment followed by proteasomal inhibition by MG132, a significant increase in the poly-Ub-XIAP pool was detected (Figure 5B). There was a substantial decrease in poly-ubiquitinated XIAP protein pool in the presence of overexpressed WT-USP7 compared to its catalytic mutant (USP7^C223S^) (Figure 5C). Furthermore, P5091 treatment was able to significantly increase XIAP polyubiquitination pool even in the presence of USP7 overexpression (Figure 5D). While detecting the nature of XIAP poly-ubiquitination, we transfected HEK cells with multiple Ub-mutants, including K-48 and K-63, to examine the poly-ubiquitination nature of XIAP. We found that USP7 efficiently deubiquitinating K-48 linked poly-ubiquitinated XIAP (Figure 5E lane 1 vs. lane 2). USP7 can also deubiquitinate K-63 linked poly-ubiquitinated XIAP, but efficiency was much lesser than K-48 linked poly-ubiquitination (Figure 5E lanes 3,4 vs. lanes 1,2). Similar results were also observed in HCT116 (p53^WT^) cells (Supplementary Figure 6A).

**Figure 5.**
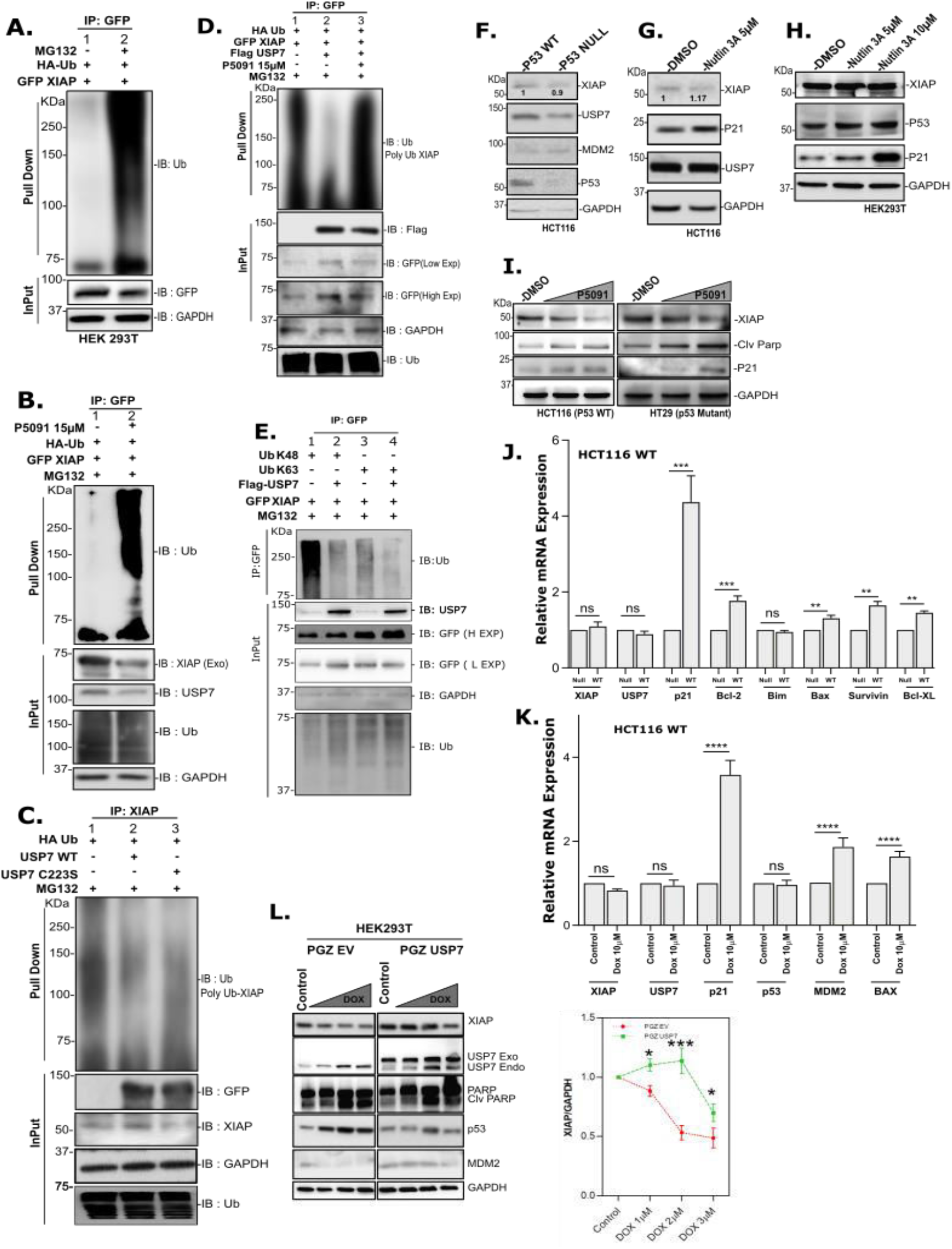
Deubiquitinase USP7 prevents ubiquitination-mediated degradation of XIAP in p53 independent fashion. HEK cells were used in experiments A-E, H & L and mentioned in other cases. **A,** Cells co-transfected with HA-Ub and GFP-XIAP and 24 hrs post-transfection cells were further treated with MG-132 (25µM) and vehicle control for an additional 4 hrs. GFP-tagged proteins were pulled down from lysates using GFP antibody and Protein-A Sepharose bead; analyzed by IB using indicated antibodies. B, Cells were transfected with indicated constructs; 24 hrs post-transfection, cells were treated with P5091 (15µM) for another 24 hrs. Before harvesting, the cells were further treated with MG-132 for 4 hrs. Lysates were prepared and used for pull-down using GFP antibody followed by IB using indicated antibodies. C, Cells were transfected individually with EV, USP7, and USP7^C223S^ plasmids. Pull-down assay was performed using XIAP antibody, bead-bound proteins and total cell lysate were analyzed by IB to detect the change in poly-ubiquitination pattern. D, Cells were transfected with indicated plasmids; 24 hr post-transfection, treated with P5091 (15µM) and vehicle control for another 24 hours. Cells were harvested after MG-132 treatment for further 4 hrs, followed by pull-down assay was performed using GFP antibody and IB with indicated antibodies. E, Cells were transfected with either WT or different Ub mutants as depicted in the figure. Following lysate preparation, pulled down the proteins using GFP antibody followed by IB using the indicated antibodies. Input (3%) was run separately for Control. F, Cell lysates prepared from p53^WT^ and p53^null^ HCT116 cells subjected to IB with the indicated antibodies. G, HCT116 (p53^wt^) cells were treated with Nutlin3A (5µM) for 24 hrs, the protein level of XIAP and other p53 responsive genes were analyzed by IB. H, Cells were treated with Nutlin3A in a dose-dependent manner (5µM and 10µM) to analyze the level of indicated proteins by IB. I, HCT116 (p53^wt^) and HT29 (p53^mut^) cells were treated with P5091 in a dose-dependent manner (0, 10, and 20µM) for 24 hrs and check the levels of the indicated proteins by IB. J, Expression of indicated genes in HCT116 cells containing either p53^wt^ or p53^mut^ was analyzed by qRT-PCR. K, HCT116 cells were treated with Doxorubicin (10µM) for 3 hrs, washed and kept in fresh media without Dox for another 21hrs before harvesting them. Expression (mRNA level) of the indicated genes was analyzed by qRT-PCR. L, Cells were transfected with either EV (left panel) or GFP-USP7 (right panel) for 24 hrs followed by Dox treatment in a dose-dependent manner (0, 1, 2, 3µM) for another 24 hrs. Expression of the indicated proteins was analyzed by IB. Expression of XIAP was normalized against GAPDH and plotted here. The data are representative of three biological replicates. qRT-PCR data represents the mean ± SD of three independent biological replicates. Indicated P-values were calculated using Student’s t-test and P<0.0001 is represented as *, otherwise non-significant (ns).

Recently it was reported that survivin, a member of the IAP family, is negatively regulated by p53^Wt^ in p53 dependent apoptosis^53^. It is to be noted that p53 and its E3 ligase MDM2 are considered as the primary targets of USP7 mediated deubiquitination in cancer. So, our next goal was to find if p53 had any role to play in the regulation of XIAP by USP7. To investigate this, we used two HCT116 cell lines (having p53^Wt^ or p53^Null^) and checked for XIAP levels. Surprisingly, we found no significant change in the XIAP protein and transcript levels (Figure 5F & 5J), where transcript and protein levels of other p53 target genes increased substantially, indicating that XIAP regulation is independent of cellular p53 status. We further justified this observation by increasing the p53 level with the help of Nutlin3A (a potent MDM2 inhibitor) in both HCT116 and HEK cells. In addition, we checked p21 protein level as a positive control for p53 activity (Figure 5G & 5H). We observed no significant change in XIAP protein level, but an increase in p53 target proteins. Additionally, we found a decrease in XIAP protein level upon USP7 inhibition by P5091 in HCT116 (p53^Wt^) and HT29 (p53^Mut^) cells (Figure 5I). Still, there is no detectable change in XIAP transcript level, although a significant increase in MDM2 transcript level was observed in HCT116 (p53^WT^) cells with respect to control (Supplementary Figure 6B). We further checked the transcript levels for USP7 and XIAP along with MDM2, p53 and p53 target genes p21, BAX in Doxorubicin (Dox) treated cells. We observed no significant change in transcripts levels of both USP7 and XIAP, although increased expression of MDM2 and p53 target genes p21 and BAX (Figure 5K). We also observed exogenously overexpressed USP7 significantly induces XIAP protein level in Dox treated HEK cells compared to the EV transfected control (Figure 5L) which may be due to USP7 mediated stabilization of XIAP. Altogether, our results suggest that USP7 deubiquitinates and stabilized XIAP in Dox-treated cells, which may result in chemoresistance against genotoxic drugs for an escape from apoptosis.

### USP7 promotes tumorigenesis and chemoresistance through XIAP stabilization

Till now, we demonstrated that XIAP is the substrate for deubiquitinase USP7 irrespective of cellular p53 status. We next questioned the physiological consequence of USP7 mediated stabilization of XIAP on tumorigenesis as XIAP is an essential endogenous inhibitor of apoptosis. For this, we have maintained a constant background level of apoptosis by treating cells with low concentration of Doxorubicin (Dox) and check for the effect of USP7 concerning USP7 mediated XIAP stabilization. HCT116 (p53^WT^) and HEK cells with elevated expression of WT-USP7 (in a dose-dependent fashion), when treated with 1µM of Dox, results in a significant increase in the levels of both endogenous and exogenous forms of XIAP protein. Consequently, there was a decrease in cleavage of PARP, caspase3 and 7, indicating that inhibition of apoptosis under genotoxic stress (Dox treatment) is due to stabilization of XIAP by USP7 (Figure 6A & 6B). This phenomenon was independent of cellular p53 status because no significant increase in p53 protein was observed. Thus, USP7 dependent stabilization of XIAP overrides doxorubicin-induced apoptosis and is independent of cellular p53 status. Furthermore, USP7 overexpression was able to increase exogenous XIAP protein level through post-translational stabilization irrespective of cellular p53 status as observed in other cancer cell lines Huh7 (p53^mut^) and HepG2 (p53^wt^) (Figure 6C).

**Figure 6.**
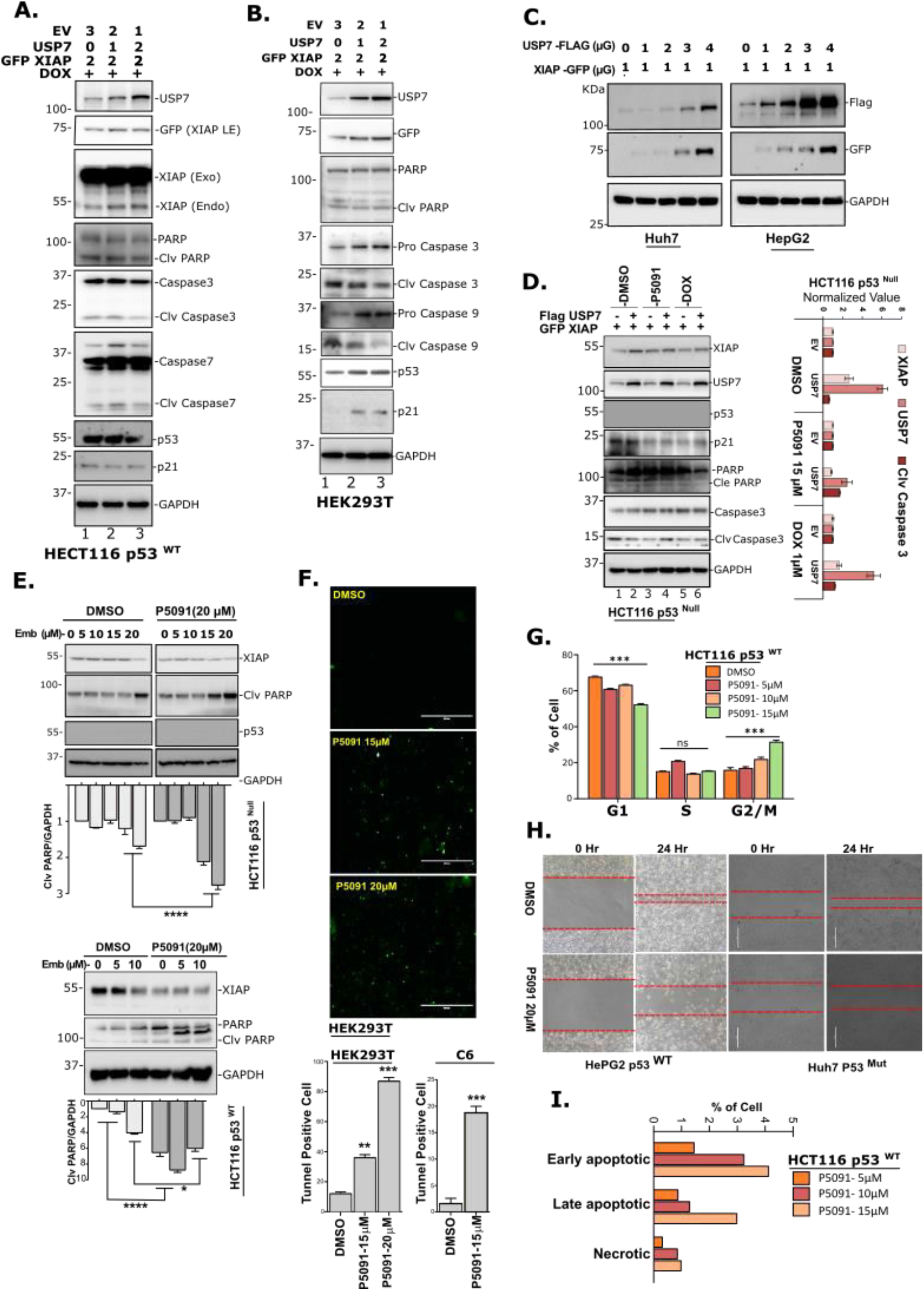
USP7 inhibits apoptosis through stabilization of XIAP. **A,** HCT 116 (p53^Wt^) cells were transfected with indicated plasmids and treated with Doxorubicin for 24 hrs to induced apoptosis, prepared the lysates and analyzed by IB using the panel of indicated antibodies. **B,** Similarly, HEK cells were transfected with indicated plasmids and treated with Doxorubicin (1 µM) for 24hrs to induced apoptosis, prepared the lysates and analyzed by IB using the panel of indicated antibodies. **C,** Huh7 (p53^Mut^) and HepG2 cells (p53^Wt^) cells were transfected with Flag-USP7 and GFP-XIAP for the indicated time points, prepared the lysates and analyzed by IB. **D,** HCT116 (p53^Null^) cells were transfected with either Flag-USP7 or GFP-XIAP. After 48 hrs, cells were treated with P5091 (15µM) or Doxorubicin (1µM). Prepared the cell lysates and analyzed by IB using the indicated antibodies. XIAP, USP7, and Cleaved Caspase 3 were plotted after normalization against GAPDH. **E,** HCT116 (p53^Null^ - upper panel) and HCT116 (p53^Wt^ - lower panel) cells were treated with either XIAP inhibitor Embellin or USP7 inhibitor P5091 for the indicated doses. Prepared the lysates and analyzed by IB. XIAP expression was plotted after normalization against GAPDH. **F,** HEK and C6 cells were seeded in a 6-well plate and treated with P5091 (15µM and 20 µM for HEK and, 15 µM for C6) for 24 hrs. Treated cells were stained as per protocol to detect TUNNEL positivity. **G,** Cell cycle profile of HCT116 (p53^WT^) cells were analyzed as per protocol after treatment with an increasing dose of P5091 (5µM, 10µM and 15µM 24 hrs). **H,** Scratch assay was performed to evaluate the migration of HepG2 (p53^Wt^) and Huh7 (p53^Mut^) cells pre-treated with P5091 (20µM). **I,** percent of cell population in different apoptosis phases of HCT116 (p53^WT^) cells were analyzed as per protocol after treatment with indicated dose of P5091 for 24 hrs. Error bars in all the indicated sub-figures represent mean (±) SD from three independent biological repeats. Indicated P-values were calculated using Student’s t-test and P<0.0001 is represented as *, otherwise non-significant (ns).

Furthermore, inhibition of apoptosis upon USP7 overexpression is reversed by P5091 mediated USP7 inhibition. Even Dox treated HCT116 (p53^Null^) cells unable to induce a significant amount of apoptosis as measured by caspase 3 cleavage upon USP7 overexpression with respect to control, which clearly indicating p53 independent anti-apoptotic function of USP7 is through stabilization of XIAP (Figure 6D). Next, we checked the magnitude of apoptosis induction by performing TUNNEL assay. There was a significant increase in cells with inter-nucleosomal DNA fragmentation with an increasing dosage of P5091. Additionally, an increase in FITC positive cells was observed with increasing dosage of P5091 with respect to DMSO control in both HEK and C6 (rat glioma) cells (Figure 6F).

According to a previously mentioned study, USP9X acts as a deubiquitinase for XIAP. The up-regulation of XIAP by USP9X promotes mitotic survival and increased resistance to mitotic spindle poisons in diffused large B-cell lymphoma^54^. Like this study, our results indicated that catalytic inhibition of USP7 with P5091 and consequent downregulation of XIAP results in G2/M phase cell cycle blockage in a dose-dependent manner in HCT116 (p53^Wt^) cells (Figure 6G). Additionally, in a scratch assay, inhibition of cell migration was observed in P5091 treated monolayer of HepG2 (p53^Wt^) and Huh7 (p53^Null^) cells (Figure 6H) supports the loss of viability and migration due to USP7 inhibition.

Next, to further support our findings, we quantified apoptotic population in P5091 treated HCT116 (p53^WT^) cells by flow cytometric analysis and found a detectable change (Figure 6I). The effect of USP7 inhibition on cellular physiology was further validated by DNA fragmentation assay where we found a noticeable increase in fragmented DNA in HCT116 and C6 cells with an increasing dose of P5091 (Supplementary Figure7A). In addition, a significant increase in late apoptotic cell number in P5091 treated HEK cells (Supplementary Figure 7B) was observed. According to a previous study, inhibition of XIAP activity by small molecule inhibitor (Embelin) results in activation of extracellular signal-regulated kinase ERK-1/2 and ROS accumulation in Inflammatory Breast Cancer^55^. Therefore, we sought to check if there is any accumulation of ROS upon USP7 inhibition. We found a dose-dependent accumulation of ROS when C6 and HCT116 cells were treated with P5091 for 24 hrs (Supplementary Figure 7D). We also detected a significant increase in Caspase activity upon dose-dependent inhibition of USP7 by either P5091 or P22077 in HCT116 cells suggesting the involvement of XIAP in the regulation of apoptosis *via* the USP7-XIAP axis (Supplementary Figure 7C).

MOPS-mediated release of pro-apoptotic Smac and the consequent inactivation of XIAP is a prerequisite for activation of apoptosis, suggesting a pro-survival effect of XIAP^56^. Another study demonstrates that XIAP can delay the mitochondrial release of Smac, Cytochrome C, and Apaf1 by inhibiting the formation of MOPS in association with the Bcl-2 family members of proteins^57^. So, we attempted to determine any effect of USP7 inhibition *via* our proposed USP7-XIAP axis on mitochondrial membrane permeabilization. Indeed, we found that with increasing dosage of USP7 inhibitor P5091, there was a significant decrease in mitochondrial membrane integrity in both HCT116 and C6 cells (Supplementary Figure 7E).

Till now, we observed that inhibition of XIAP with its small molecule inhibitor Embelin additionally supports P5091 mediated apoptosis induction in both p53^Wt^ and p53^Null^ HCT116 cells (Figure 6E upper and lower panels) as evidenced by an increase in PARP cleavage and decrease in XIAP level as shown in the immunoblots. Therefore, we can conclude that combinatorial inhibition of USP7 and XIAP can induce apoptosis in a higher magnitude than their individual inhibition.

### Therapeutic efficacy of nanoformulated USP7 inhibitors *in vitro*

To examine and validate our findings, we developed nanotized USP7 inhibitors, P5091 and P22077 as PLGA-based nanoformulations to effectively internalize the inhibitors in cells even at low concentrations. Figure 7 shows the characterizations and optimization studies of USP7 inhibitors packed nanoformulations. The encapsulation efficiencies of P5091 and P22077 nanoparticles are 95% and 85% respectively at an optimum drug-polymer ratio of 1:5. Lyophilized powder of these nanoformulations are stable, readily dissolved in water, and could be stored at room temperature and at 4°C without any decomposition or aggregation. Transmission electron microscopy (TEM) (Figure 7A) and atomic force microscopy (AFM) (Figure 7B) was performed with an aqueous dispersion of these nanoparticles, showed their spherical shape and size of the nanoparticles from 40 to 75 nm. Zeta potential values of empty, P5091, and P22077 nanoparticles ranges from 12 to 50 mV (Figure 7C), respectively. The values imply higher stability of all the nanoformulations. Additionally, no structural changes of these nanoformulations was confirmed by Fourier Transform Infrared Spectroscopy (FTIR) (Supplementary Figure S8B).

**Figure 7.**
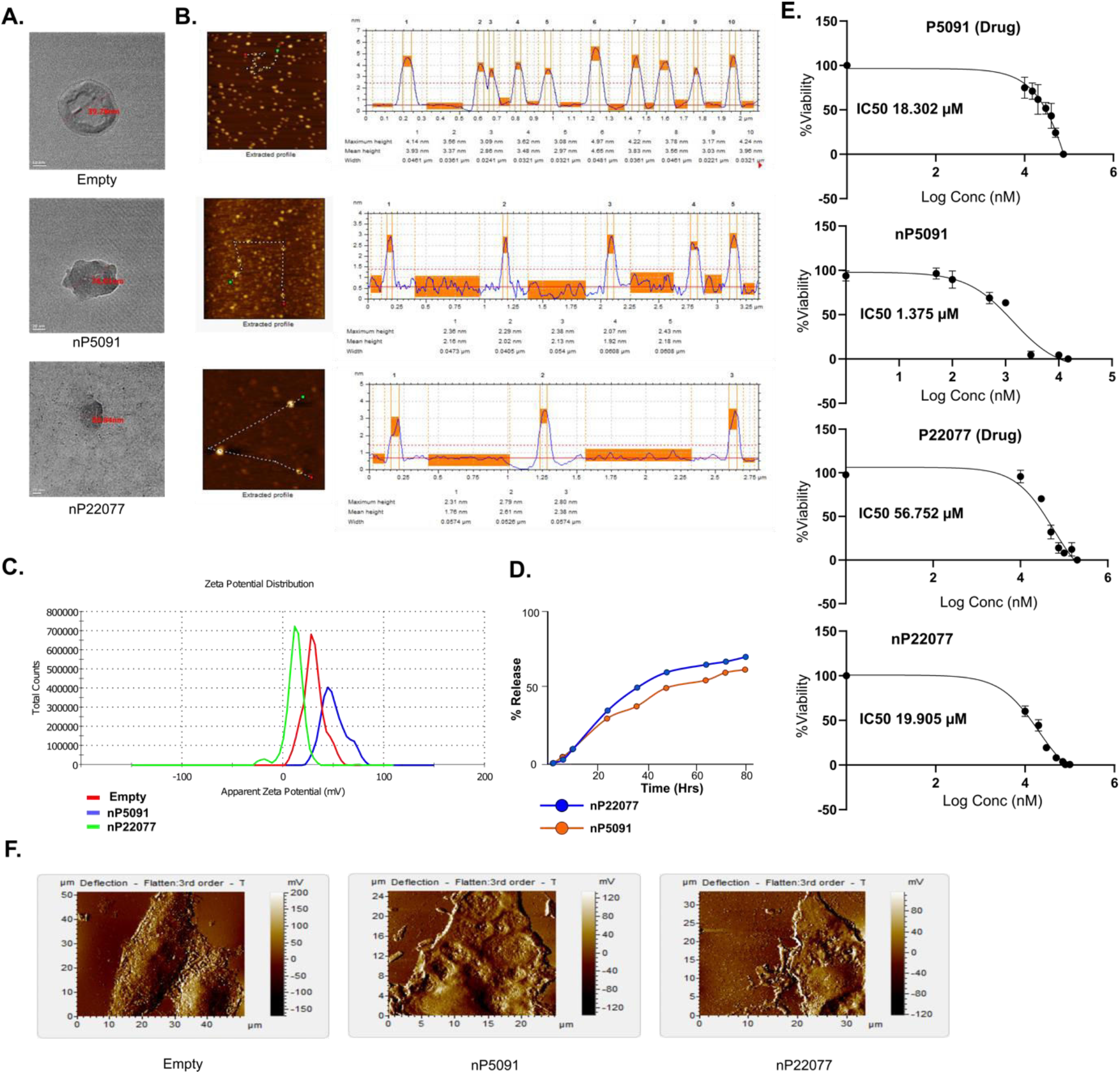
Therapeutic efficacy of nanoformulated USP7 inhibitors *in vitro*. **A,** TEM images of empty, nP5091 and nP22077 nanoparticles. **B,** AFM images of empty, nP5091 and nP22077 nanoparticles showing particle size ranging from 40 to 60nm. **C,** Zeta potential values of empty, nP5091 and nP22077 nanoparticles. **D,** Kinetic release of P5091 and P22077 from their respective nanoformulations were shown. **E,** *In vitro* cytotoxicity study of rat glioma C6 cells treated with P5091 and P22077 and their respective nanoformulations. **F,** Changes in cellular architecture of C6 cells treated with either empty, nP5091, or nP22077 nanoformulation.

Furthermore, their thermal stability has also been validated by TGdTA (Supplementary Figure 8A) analysis which indicates that nanotized compounds were stable at higher temperatures. The comparative thermal profile of PLGA nanoparticles exhibited two sharp exothermic peaks that confirmed thermal decomposition of amorphous polymer vesicles. Thermal decomposition of nanoformulated P5091 and P22077 begun at the same temperature as the empty nanoparticles, but the exothermic event was more intense because of decomposition. From all these characterizations, it has been suggested that positive surface charges of DMAB helped in the successful entrapment of the compounds in PLGA nanoparticles. The residual aggregation of the nanoparticles if any due to their charged surface was also controlled by sonication before experimental use. Furthermore, PLGA nanoparticles released P5091 and P22077 in a controlled and efficient manner with a small percentage of drugs retained in the nanovesicles after 24 hours (Figure 7D). This implied the robust encapsulation of both the compounds.

Our major concern was to ascertain any cytotoxicity for utilization of FDA approved PLGA-based nanoformulation as drug delivery vehicles for cancer therapy. Hence, cytotoxicity of nanoformulated compounds as well as their native forms were evaluated by MTT assay (Figure 7E). The viability of rat glioma C6 cells after 24 hrs of treatment with USP7 inhibitors P5091 and P22077 was determined. We observed that USP7 inhibitor P22077 has less efficiency when compared with P5091 in both respective nanoformulations and their native forms (IC-50 values are P5091:18.302µM, nP5091:1.375µM; P22077:56.752µM and nP22077:19.905µM). Our AFM result of nanoformulations treated C6 cells indicates that nP5091 caused more apoptotic effects like cellular deformation even at lower concentration than nP22077 within 24 hrs (Figure 7F) indicating P5091 has better effectivity than nP22077.

## Discussion

Apoptosis is essential for the removal of cells that have gathered irreparable genomic damage and is therefore crucial for preventing the propagation of cells whose genomic integrity has been compromised^58^. A better understanding of the underlying mechanisms regulating apoptosis is very important as it plays a vital role in the pathogenesis of several diseases. For example, several degenerative diseases are associated with excessive apoptosis and tissue degeneration, whereas diseases like cancer show suppression of apoptosis resulting in the generation of immortalized malignant cells. Furthermore, apoptosis is a complex and multistep process, and so defects at any point may result in malignant transformation of affected cells and development of drug resistance.

Aberrant expressions of Inhibitor of Apoptosis Proteins (IAPs) are associated with many cancers. For example, it was demonstrated that overexpression of cIAP-2 in pancreatic cancer was responsible for the development of chemoresistance^59^. Similarly, another member of the IAP family, Apollon was upregulated in glioma and responsible for cisplatin and camptothecin resistance^60^. Survivin and XIAP were also found to be upregulated in non-small cell lung carcinoma (NSCLC). The study revealed that over-expression was associated with resistance to a variety of apoptosis-inducing anti-cancer agents^61^.

Several studies into the functional aspects of USP7 have revealed that the primary mechanism underlying the induction of apoptosis upon USP7 inhibition is the restoration of the tumor suppressor p53^62^. However, the mechanism responsible for facilitating the same effect upon USP7 inhibition in p53^Null^ cells has not received much attention previously. We observed that apoptosis induction and growth retardation due to USP7 inhibition is similar in cell lines with varied p53 status like wild-type, mutant and even null background. Moreover, cell viability and apoptosis studies carried out on USP7 inhibitor-treated p53^Wt^ or p53^Null^ HCT116 cells indicated that p53 is not a solo participant in USP7 inhibition-mediated apoptosis induction. Thus, there may be some unexplored, alternative signaling pathways that are under the control of USP7 (Figure 8). To address this issue, we have used a proteomics-based approach and identified some novel USP7 interacting proteins, of which some molecules were important in the regulation of apoptosis. After thorough analysis, we successfully identified the inhibitor of apoptosis protein XIAP as a novel interactor of USP7. In fact, we have identified a positive correlation between USP7 and XIAP expressions in an array of cancer cell lines belonging to multiple cancer types, which clearly indicates the importance of USP7 and XIAP in cancer progression and preventing apoptosis in cancer therapy.

**Figure 8.**
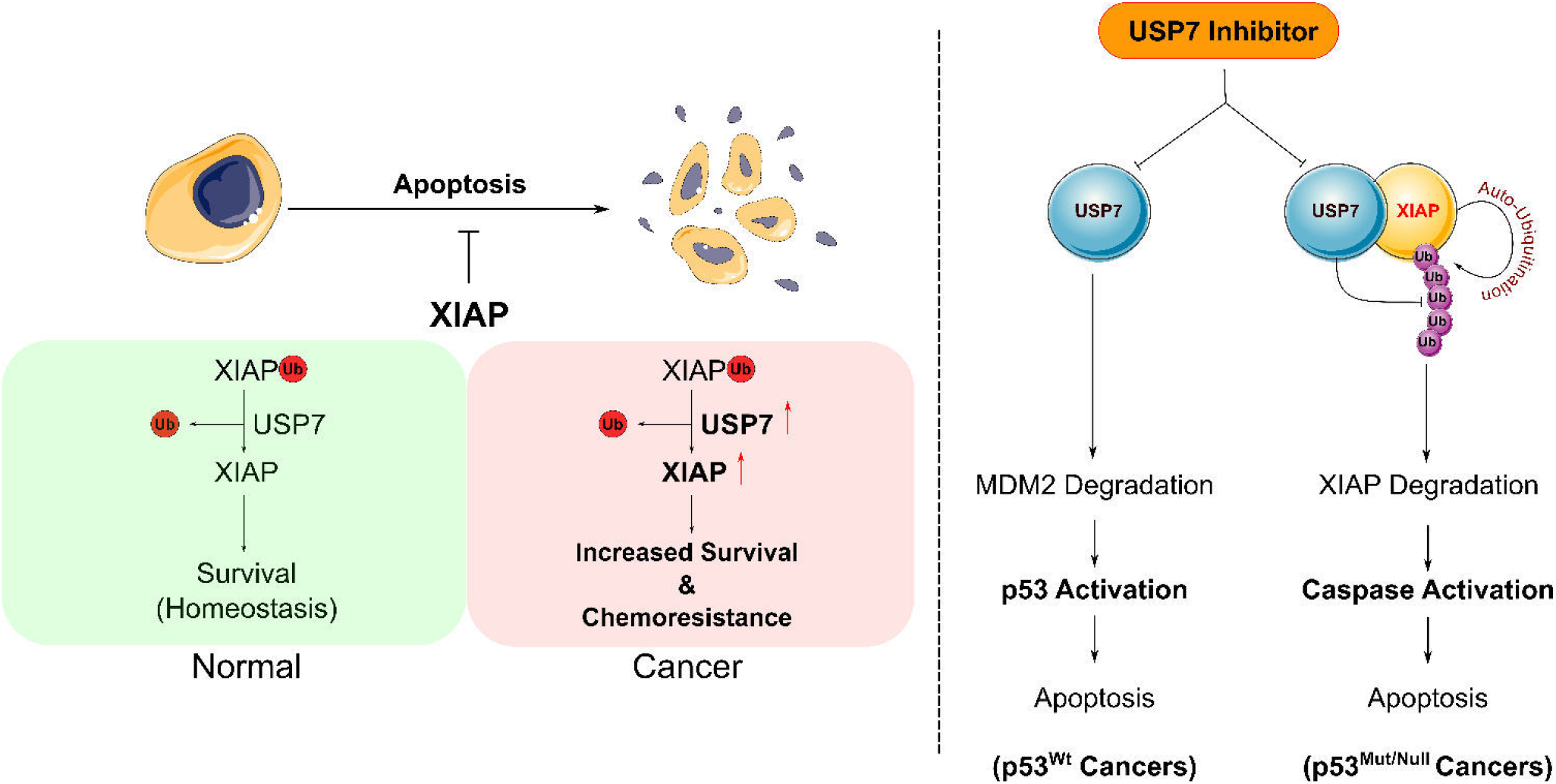
Proposed model. USP7 deubiquitinates and stabilizes anti-apoptotic protein XIAP and promote cancer cell survival. Overexpression of either USP7 or XIAP confers chemotherapy resistance and associated with increased survival of cancer cells. USP7 inhibition by small molecule inhibitor acts as a potential therapeutic intervention to fight against both p53^Wt^ and p53^Mut/Null^ cancers through MDM2 and XIAP respectively.

In this report, through modulation of USP7 expression and functional inhibition we demonstrated that USP7 regulates XIAP stability in both normal and cancer cell lines, and the deubiquitinase activity of USP7 is essential for XIAP function. We also establish that despite its natural tendency to sequester to the nucleus, USP7 can inhabit the cytoplasm where it colocalizes with XIAP. By interaction-based studies, we can strongly conclude that USP7 does indeed physically interact with XIAP through a conjunction of its UBL repeats and the catalytic domain. By developing an array of domain deletion mutants for XIAP, we showed that the BIR2 and the linker region between BIR2 and BIR3 are necessary for interaction with USP7. A marked decrease or increase in the poly-ubiquitination pattern for XIAP was observed upon over-expression or depletion of cellular USP7 protein levels, respectively. One of the well-established mechanisms by which p53 induces apoptosis is through the transcriptional down-regulation of inhibitors of apoptosis factors such as survivin. Additionally, we determined that p53 does not exert any transcriptional control over *XIAP* gene. The observed deubiquitination and consequent stabilization of XIAP by USP7 is a completely p53-independent process. Our results indicate that stabilization of XIAP by USP7 leads to the overall development of an anti-apoptotic state within the cell. USP7 mediated stabilization of XIAP counteracts the induction of PARP cleavage and activation of Caspases by apoptotic stimuli. Additionally, stabilized XIAP also led to reduced accumulation of intracellular ROS and aided in the maintenance of mitochondrial membrane integrity, thereby impeding the release of pro-apoptotic factors like Smac, Cytochrome c, and Apaf1, which agrees with the previous studies^55–57^.

In our study, treatment of cancer cells with an optimized PLGA-based nanoencapsulated USP7 inhibitors P5091 and P22077 have improved solubilities and anti-cancer activities at much lower concentrations compared to their native forms. Physicochemical characterizations performed with nP5091 and nP22077 clearly indicate that size of the nanoparticles were within the range of 40-75 nm. These nanoformulations can readily dissolved in water and zeta potential values indicate that these nanoparticles exert higher stability. Moreover, FTIR analysis concluded that P5091 and P22077 have no structural changes in nanoformulations. These nanoparticles are also biocompatible and less toxic. The presence of DMAB played a critical role in nanoparticle formation with more entrapment efficiency in comparison to loading efficiency. Previously, we showed that formation of nanoparticles with DMAB exerted significant protective effects against cellular disorder^63^. Moreover, it was used as a stabilizer leading to dispersion of positive charges on nanoparticles. It can easily associate with the cells through electrostatic interactions and releasing the drugs in a more efficient manner at the target site. It also exhibited perinuclear localization of the compounds followed by endosomal escape. AFM studies has been clearly shown that nanocompounds acted on rat glioma C6 cells and diffused across the nuclear membrane into the nucleus thereby resulting in severe damage of organelles results into change in cellular architectures with onset of apoptosis. Cytotoxicity study indicates that nanoformulated P5091 and P22077 (nP5091 and nP22077) showed higher potency for killing C6 cells in comparison to normal drugs. These results clearly signifies that nanoformulated compounds had higher cellular uptake and sustained release behaviour.

In conclusion, in this study, we have identified E3 ubiquitin ligase XIAP as a novel substrate for deubiquitinase USP7. The stabilization of XIAP by USP7 mediated deubiquitination is a p53-independent process. Furthermore, XIAP exhibits potent anti-apoptotic effects within the cell. The degradation of XIAP represents a novel and p53-independent mechanism of apoptosis induction upon USP7 inhibition. Further, in this study, we observed that combinatorial inhibition of USP7 and XIAP can induce cellular apoptosis in a higher magnitude than their individual inhibition. We have selected USP7 inhibitor P5091 and its nanoformulation alone or in combination with XIAP inhibitor (Embelin) to explore further in this new axis of cancer therapy.

## Supporting information

BioRxiv_Supplementary file_Ghosh MK

## Author Contributions

GS and MKG conceptualized the study, designed and analyzed the data. GS, performed all the experiments, data acquisition, and statistical analyses. SS and PSM synthesized, characterized the nanoformulations and performed *in vitro* cell culture-based experiments. KK performed all the bioinformatic analyses under the guidance of SC. GS, KK and MKG wrote the manuscript.

## Data Availability

Data supporting the conclusion of the current study is included in the manuscript and the supplementary files.

## Acknowledgements

We sincerely thank to Mr. Bhaskar Basu and Mrs. Srija Roy, (CSIR-IICB) for technical assistance. In addition, we also thank to Dr. Malini Basu and Mr. Bhaskar Basu for editing the manuscript.

## Supplementary data

Supplementary figures S1 – S8; List of cloning primers (A), List of Real-Time PCR (RT-PCR) primers (B); and Supplementary tables S1 - S4.

