## Supplementary material for "Identification and validation of E3 ubiquitin ligase XIAP as a novel substrate of deubiquitinase USP7 (HAUSP) - Implication towards oncogenesis": BioRxiv_Supplementary file_Ghosh MK

### Supplementary figures (S1-S8)

#### Figure S1

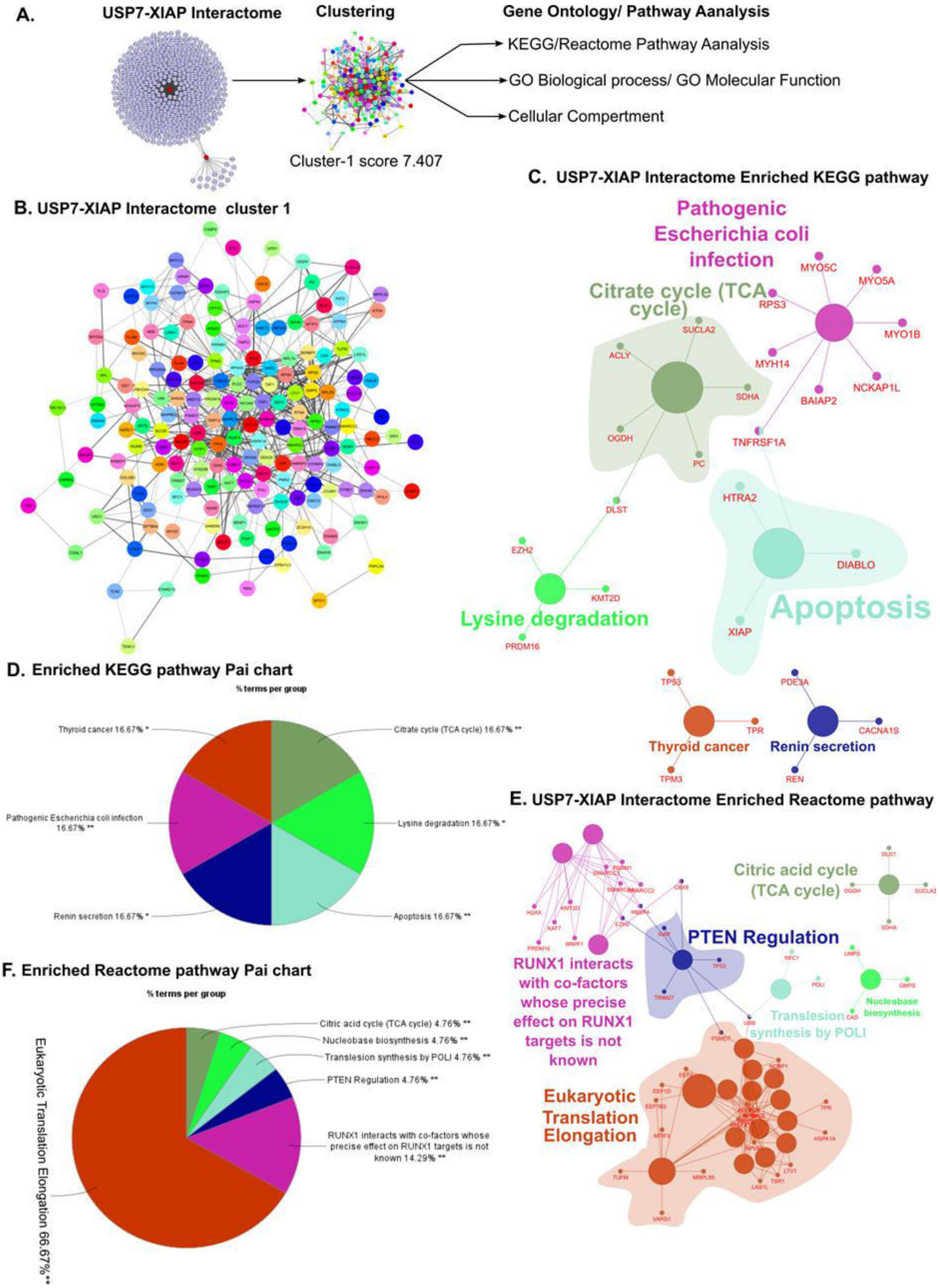

#### Figure S2

**A.** USP7-XIAP Interactome Enriched GO Biological Process

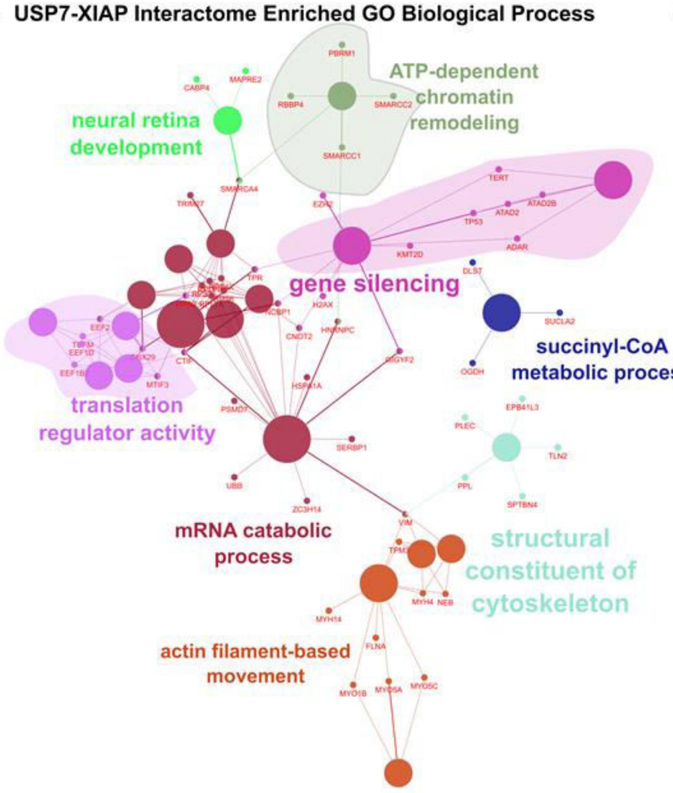


**B.** Enriched GO Biological Process Pai chart

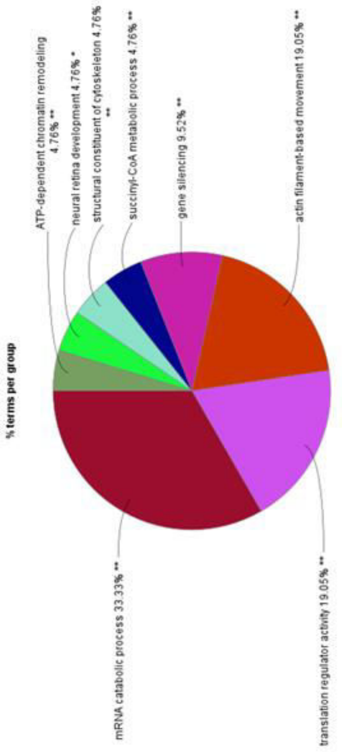


**C.** USP7-XIAP Interactome Enriched GO Molecular Function

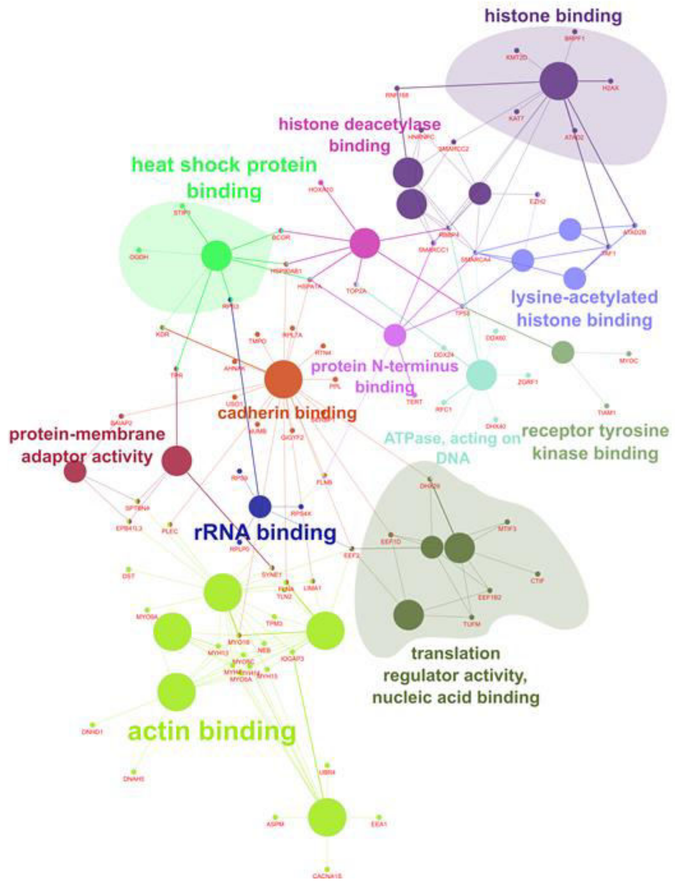


**D.** Enriched GO Biological Process Pai chart

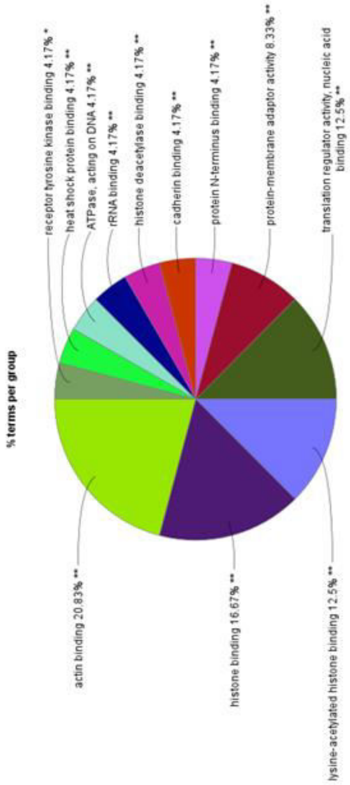

#### Figure S3

A.

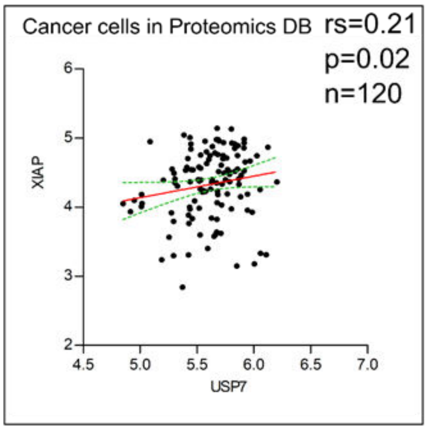

Cancer cells in Proteomics DB
rs=0.21
p=0.02
n=120
XIAP
USP7


B.

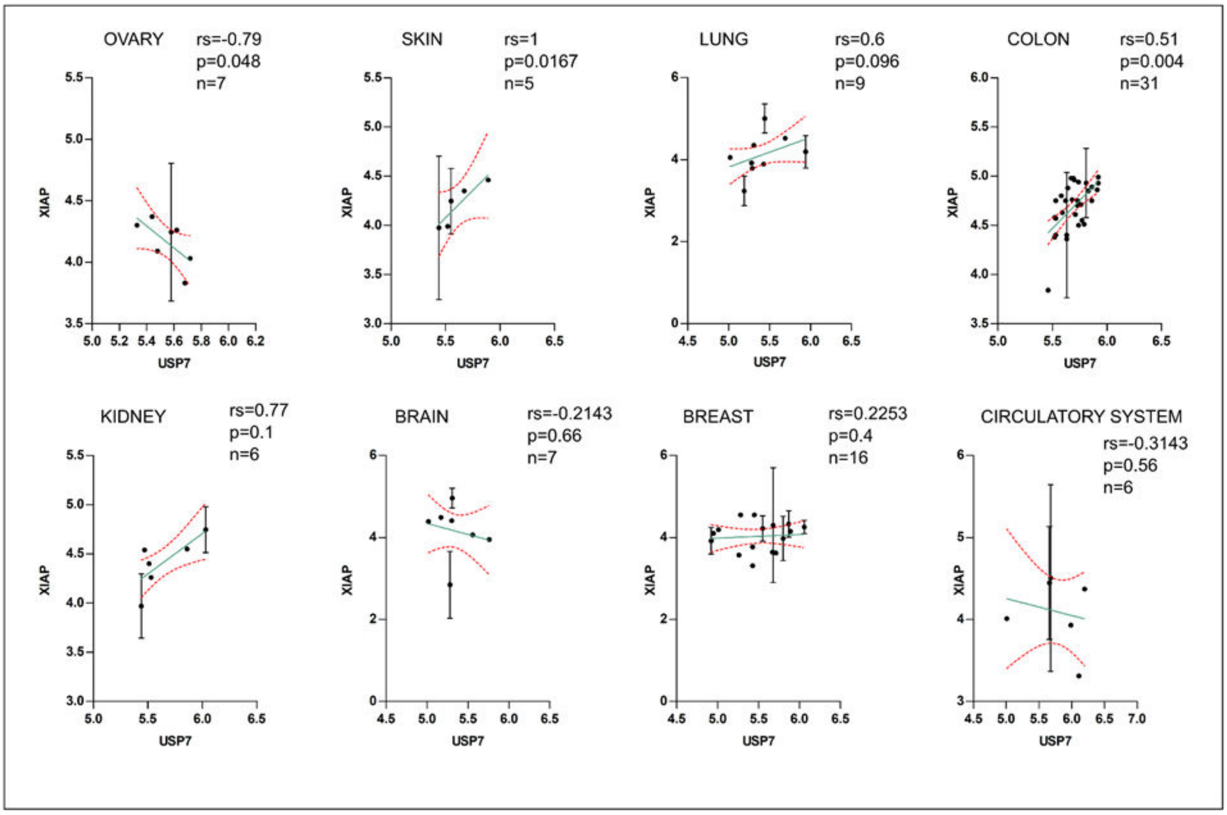

OVARY
rs=-0.79
p=0.048
n=7
SKIN
rs=1
p=0.0167
n=5
LUNG
rs=0.6
p=0.096
n=9
COLON
rs=0.51
p=0.004
n=31
KIDNEY
rs=0.77
p=0.1
n=6
BRAIN
rs=-0.2143
p=0.66
n=7
BREAST
rs=0.2253
p=0.4
n=16
CIRCULATORY SYSTEM
rs=-0.3143
p=0.56
n=6
XIAP
USP7

### Figure S4

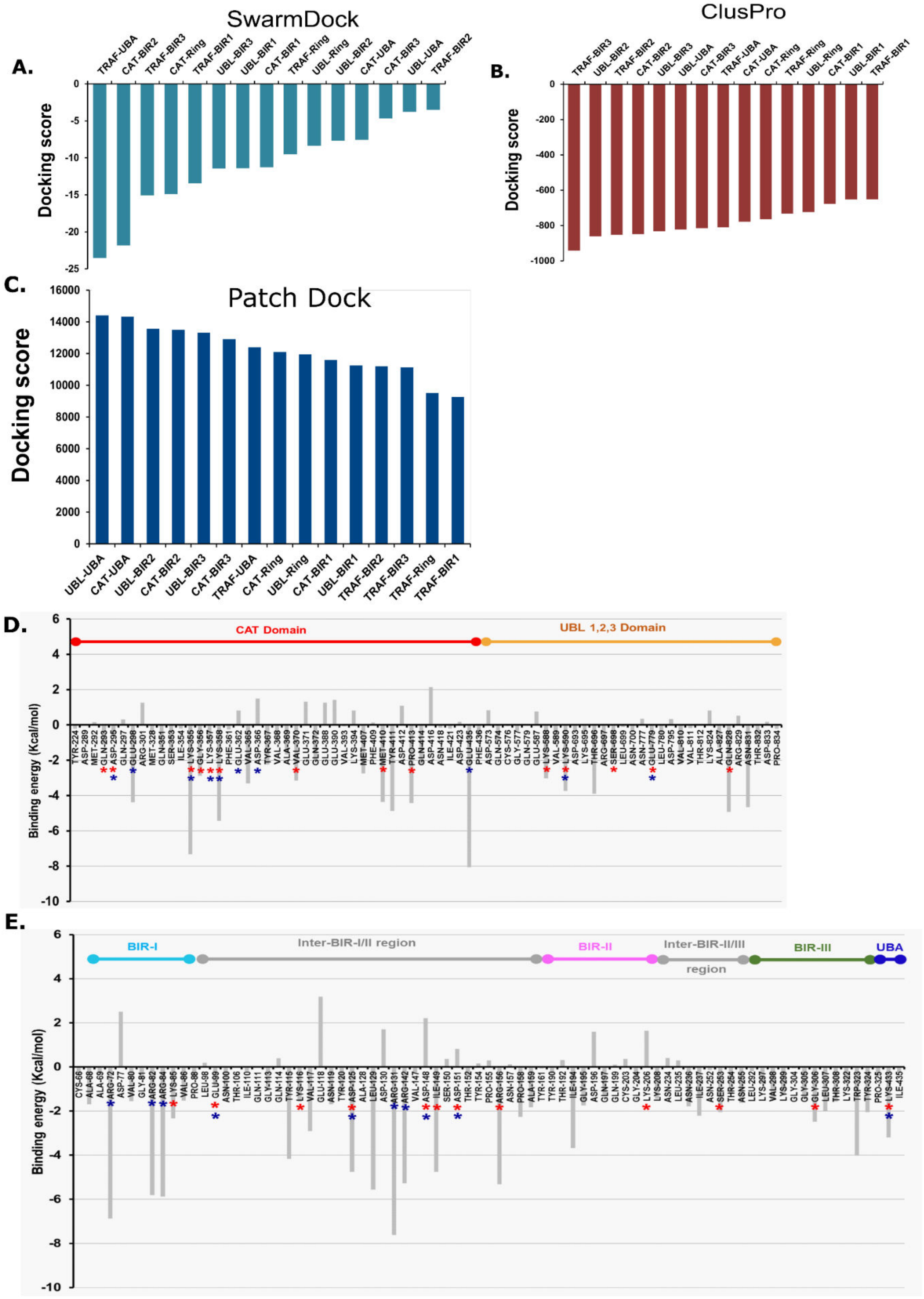

A.
SwarmDock
Docking score
TRAF-UBA
CAT-BIR2
TRAF-BIR3
CAT-Ring
TRAF-BIR1
UBL-BIR3
UBL-BIR1
CAT-BIR1
TRAF-Ring
UBL-Ring
UBL-BIR2
CAT-UBA
CAT-BIR3
UBL-UBA
TRAF-BIR2
0
-5
-10
-15
-20
-25
B.
ClusPro
Docking score
TRAF-BIR3
UBL-BIR2
TRAF-BIR2
CAT-BIR2
UBL-BIR3
UBL-UBA
CAT-BIR3
TRAF-UBA
CAT-UBA
CAT-Ring
TRAF-Ring
UBL-Ring
CAT-BIR1
UBL-BIR1
TRAF-BIR1
0
-200
-400
-600
-800
-1000
C.
Patch Dock
Docking score
16000
14000
12000
10000
8000
6000
4000
2000
0
UBL-UBA
CAT-UBA
UBL-BIR2
CAT-BIR2
UBL-BIR3
CAT-BIR3
TRAF-UBA
CAT-Ring
UBL-Ring
CAT-BIR1
UBL-BIR1
TRAF-BIR2
TRAF-BIR3
TRAF-Ring
TRAF-BIR1
D.
CAT Domain
UBL 1,2,3 Domain
Binding energy (Kcal/mol)
6
4
2
0
-2
-4
-6
-8
-10
E.
BIR-I
Inter-BIR-I/II region
BIR-II
Inter-BIR-II/III region
BIR-III
UBA
Binding energy (Kcal/mol)
6
4
2
0
-2
-4
-6
-8
-10

### Figure S5

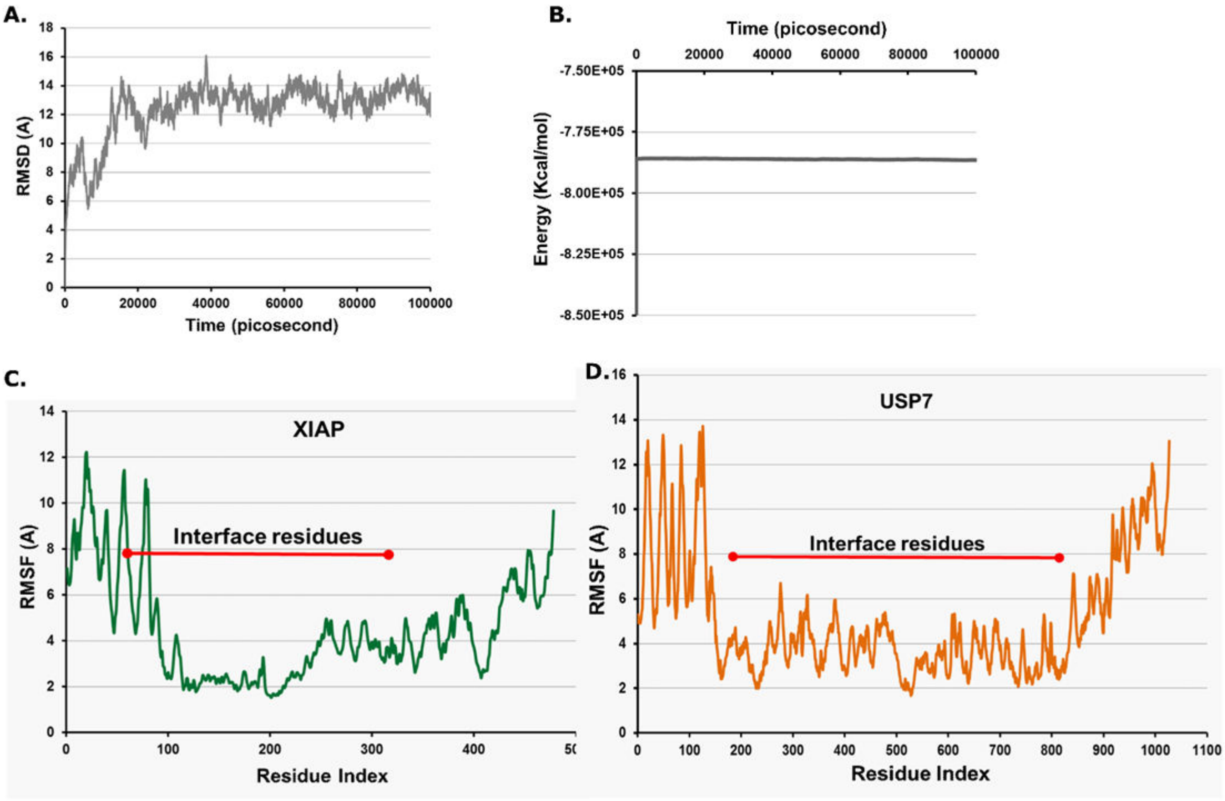

A.
RMSD (A)
Time (picosecond)
B.
Time (picosecond)
Energy (Kcal/mol)
C.
XIAP
Interface residues
RMSF (A)
Residue Index
D.
USP7
Interface residues
RMSF (A)
Residue Index

### Figure S6

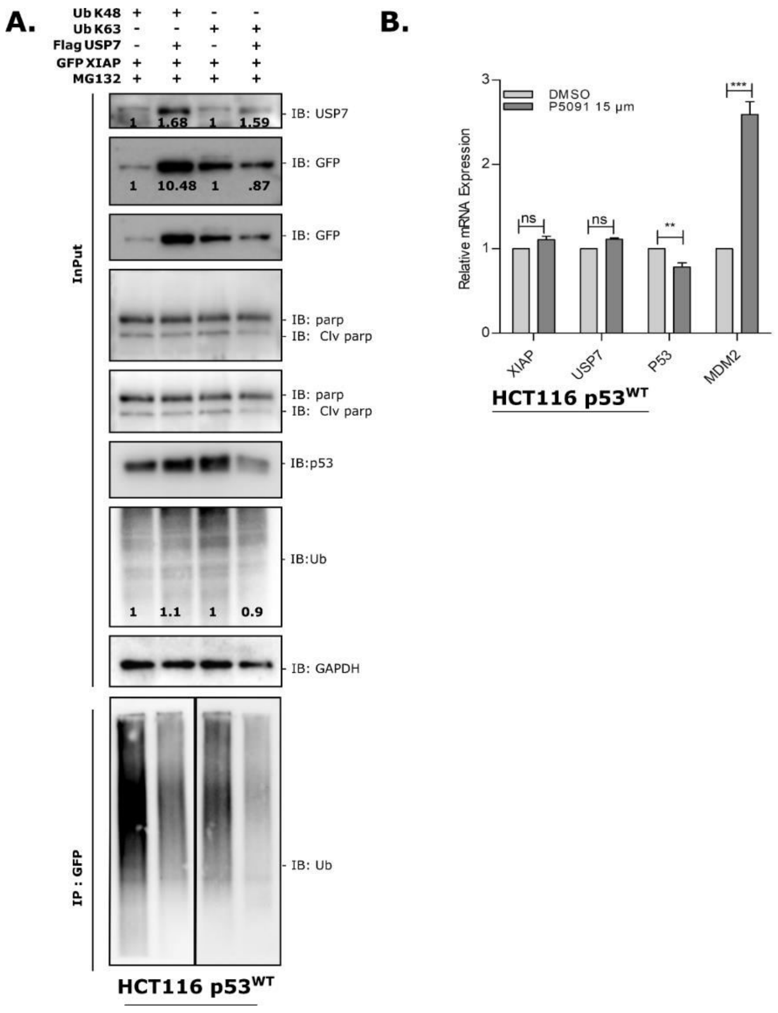
A.
Ub K48 + + - -
Ub K63 - - + +
Flag USP7 - + - +
GFP XIAP + + + +
MG132 + + + +
1 1.68 1 1.59
IB: USP7
1 10.48 1 .87
IB: GFP
IB: GFP
IB: parp
IB: Clv parp
IB: parp
IB: Clv parp
IB:p53
1 1.1 1 0.9
IB:Ub
IB: GAPDH
InPut
IP : GFP
IB: Ub
HCT116 p53WT
B.
DMSO
P5091 15 µm
Relative mRNA Expression
ns
ns
**
***
XIAP
USP7
P53
MDM2
HCT116 p53WT

#### Figure S7

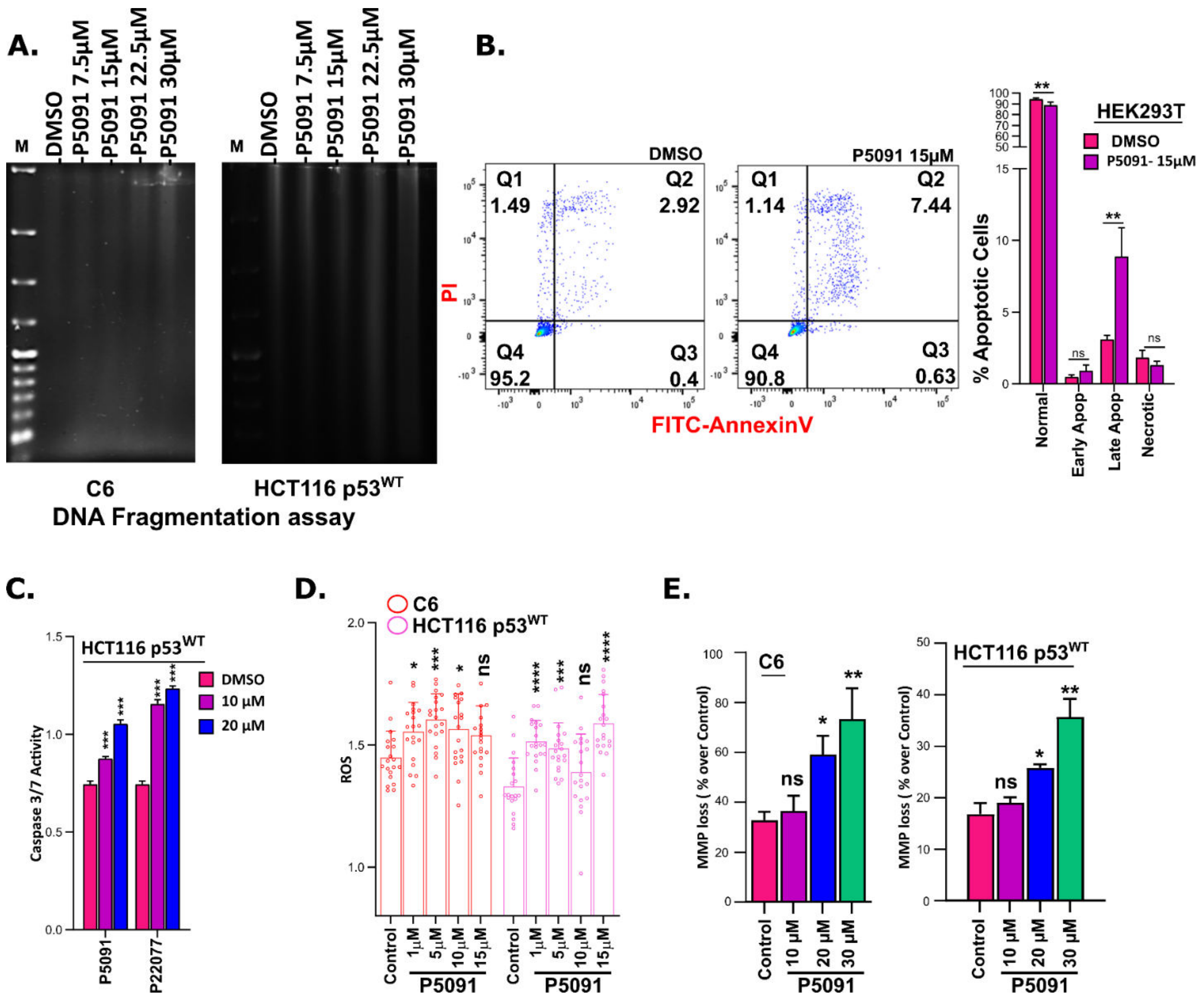
A.
M
DMSO
P5091 7.5µM
P5091 15µM
P5091 22.5µM
P5091 30µM
C6
HCT116 p53WT
DNA Fragmentation assay
B.
DMSO
P5091 15µM
Q1 1.49
Q2 2.92
Q4 95.2
Q3 0.4
Q1 1.14
Q2 7.44
Q4 90.8
Q3 0.63
PI
FITC-AnnexinV
HEK293T
DMSO
P5091- 15µM
% Apoptotic Cells
Normal
Early Apop
Late Apop
Necrotic
C.
HCT116 p53WT
DMSO
10 µM
20 µM
Caspase 3/7 Activity
P5091
P22077
D.
C6
HCT116 p53WT
ROS
Control
1µM
5µM
10µM
15µM
P5091
E.
C6
HCT116 p53WT
MMP loss ( % over Control)
Control
10 µM
20 µM
30 µM
P5091

#### Figure S8

A. B.

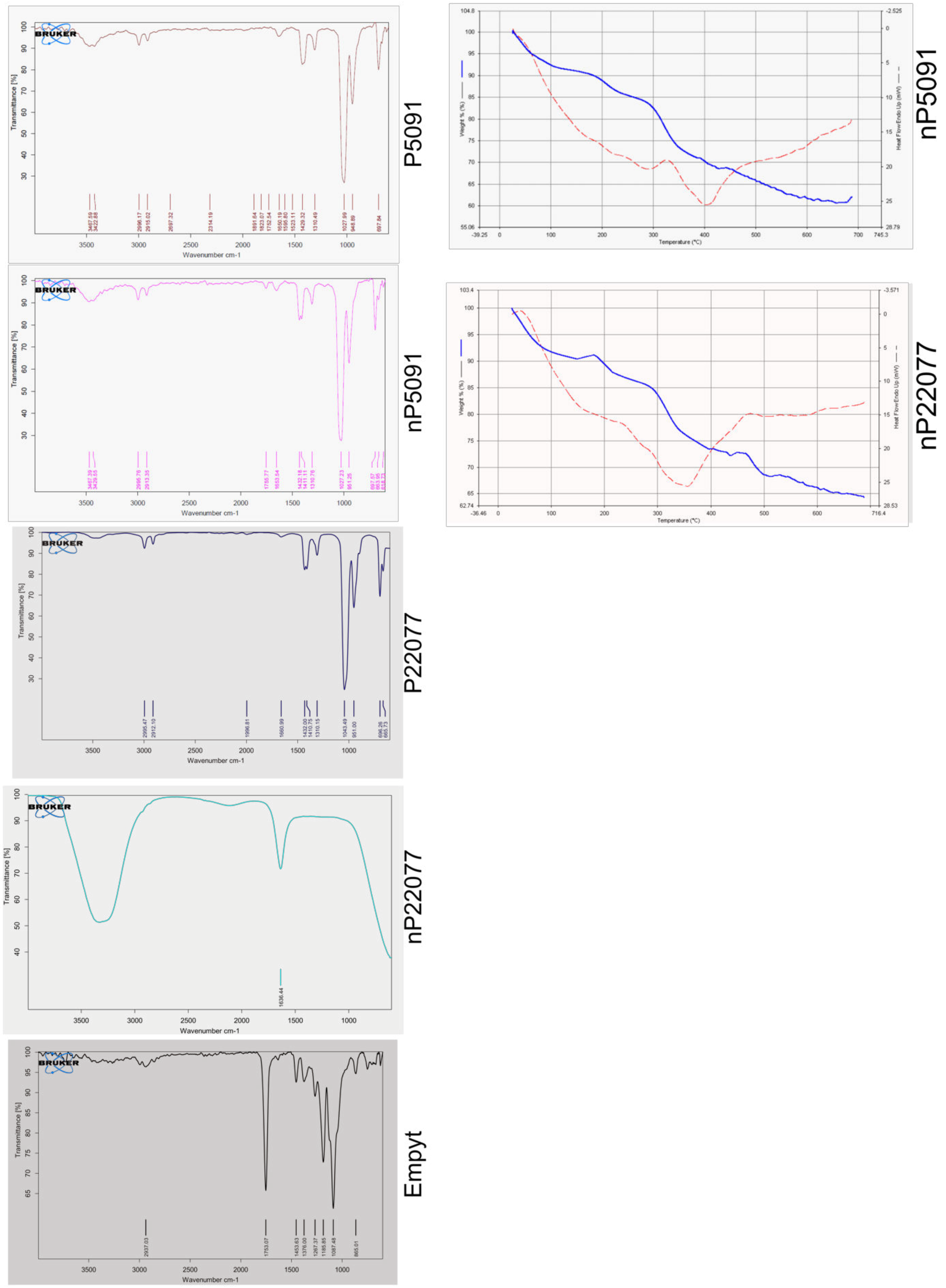
P5091
nP5091
P22077
nP22077
Empyt
BRUKER
Transmittance [%]
Wavenumber cm-1
Weight % (%)
Heat Flow Endo Up (mW)
Temperature (°C)
nP5091
nP22077

### Supplementary Figure Legends

#### Supplementary figures (S1-S8)

##### Supplementary figure S1:

A. The PPI network formed by USP7-XIAP interacting proteins in String Network analyzer of Cytoscape application with minimum interaction score of 0.4. The hub gene network was analyzed by MCODE for the PPI network. Degree≥15, MCODE scores > 5, degree cut-off = 2, node score cut-off = 0.2, max depth = 100 and k-score = 2. The largest connected cluster with the score of 7.407 was further analyzed for Gene Ontology and enriched pathways in the ClueGO Cytoscape plugin.

B. Largest hub gene network analyzed in MCODE from USP7-XIAP interactome.

C. USP7-XIAP interactome enriched with KEGG pathway terms. Circle size and colour represent the number of genes involved and specific KEGG pathway terms respectively. Involved genes enriched in the KEGG pathway belong to the USP7-XIAP interactome denoted in red.

D. Distribution of functional categories of KEGG pathway terms associated with USP7-XIAP interactome. The size of each type within the pie chart represents the percentage of included terms. Enriched KEGG terms, which are statistically significant denoted single (*) or double (**) asterisk, indicate significant enriched KEGG terms at the $p < 0.05$ and $p < 0.01$ statistical levels, respectively.

E. Distribution of functional categories in Reactome pathway terms enriched in cluster 1 of USP7-XIAP interactome. Different enriched pathway terms are coloured differently, with essential connector genes represented in red.

F. The percentage of included terms and their statistical significance from the Reactome pathway enrichment analysis is represented in the Pai chart, where statistically significant denoted single (*) or double (**) asterisk indicates significantly enriched Reactome pathway terms at the $p < 0.05$ and $p < 0.01$ statistical levels, respectively.

##### Supplementary figure S2

A - D. The similar process is repeated for GO Biological process and GO Molecular Functions for Cluster 1, as mentioned in supplementary figure 1, using MCODE and ClueGO plugins in Cytoscape.

#### Supplementary figure S3

A. Correlation analysis of USP7 and XIAP normalized ms1 peptide intensities from 120 different cancer cell lines belongs to different cancer regimes showing a positive correlation ($r_s$= 0.21) with a significant p-value of 0.02.

B. Correlation analysis of USP7 and XIAP normalized ms1 peptide intensities from cells lines in different cancer types showing both positive and negative correlations. Type of correlation, number of samples, $r_s$, and p values mentioned in the figure.

#### Supplementary figure S4

A-C. The USP7-XIAP domain pairs from each program were ranked based on the average docking score of the solutions in the largest cluster. Docking score of domain-wise docking of USP7 and XIAP obtained from three independent docking software, i.e., Swarmdock, Claspro, and Patch dock. The top six USP7-XIAP pairs were UBL-UBA, CAT-UBA, UBL-BIR2, CAT-BIR2, UBL-BIR3, and CAT-BIR3 when docking was performed using PatchDock. TRAF-BIR3, UBL-BIR2, TRAF-BIR2, CAT-BIR2, UBL-BIR3, and UBL-UBA were the top six USP7-XIAP pairs when docking was performed using ClusPro docking program. SwarmDock resulted in TRAF-UBA, CAT-BIR2, TRAF-BIR3, CAT-Ring, TRAF-BIR1, and UBL-BIR3 top six USP7-XIAP pairs.

D&E. Interface amino acids of USP7 and XIAP complex and their binding energy regarding the capacity of salt bridge formation.

#### Supplementary figure S5

A. The selected USP7-XIAP complex was subjected to MD simulation for 100 ns. As a result, favorable structural adaptation in the USP7-XIAP complex was observed, and the complex getting more stable during the simulation as detected by an increase in mean RMSD value.

B. Data represent the maintenance of a stable energy state throughout the simulation run of 100 ns, suggesting the stability of the complex during simulation.

C. The most negligible fluctuation of interfacing residue of USP7 and XIAP suggesting better stability of the complex (C, D).

#### Supplementary figure S6

A. HCT116 $p53^{WT}$ cell line was transfected with different Ub mutants (K48, K63). Following lysate preparation, pull down the proteins using the GFP antibody. Ubiquitination was detected by IB using the indicated antibodies. 3% input was run separately for control.
B. Expression of indicated genes in HCT116 $p53^{WT}$ cells treated with 15μM of USP7 inhibitor P5091 for 24 hrs was analyzed by qRT-PCR. Data represent the mean ±SD of three independent biological replicates.

#### Supplementary figure S7

A. The amount of DNA fragmentation due to apoptosis was detected in HCT116 $p53^{WT}$ and C6 cell lines treated with the indicated concentration of P5091 for 24 hrs. In addition, DNA was isolated and ran in 0.8% agarose gel and stained with EtBr for visualization.
B. FACS analysis (Annexin V /PI) of HEK cells to determine cell population at different apoptotic phases. Cells were treated with 15uM USP7 inhibitor P5091 and collected after 24 Hrs. Treated cells were further prepared for Annexin V, PI staining as per manufacturer protocol. Data represented mean ± SD of two independent biological replicates. A significant change with respected control denoted by astricts (*) where nonsignificant value as ns.
C. Caspase 3/7 activity was measured in HCT116 $p53^{WT}$ treated with 10 μM and 20μM of USP7 inhibitor P5091 and P22077. Data represented mean ± SD of three independent biological replicates.
D. Reactive oxygen species (ROS) generation was measured in C6 and HCT116 $p53^{WT}$ cells treated with indicated concentrations of P5091 for 24 hrs. Data representative of three biological replicates, each with three technical replicates as Mean ± SD.
E. Mitochondrial membrane potential loss over control was measured in C6 and HCT116 ($p53^{WT}$) cells with the help of JC-10 upon treatment with indicated concentrations of P5091 for 24 hrs. Data representative of three independent biological replicates, Mean ± SD.

#### Supplementary figure S8

Physical characterizations of empty nanoparticles, nP5091 and nP22077.

A. FTIR spectra of P5091, P22077 and their nanoformulations. Additionally, FTIR spectra of empty nanoparticles was also done for comparison.
B. TG/DTA thermograms for nP5091 and nP22077 particles.

### Additional file - 1

#### *A. List of cloning primers used*

| | |
|---|---|
| USP7_F | 5’ – AATGGATCCATGAACCACCAGCAGCAGCAGCAG – 3’ |
| USP7_R | 5’ – AATCTCGAGGTTATGGATTTTAATGGCCTTTTCAAG – 3’ |
| USP7-207_R | 5’ – AATCTCGAGTGAATCCCACGCAACTCCATGGGG – 3’ |
| USP7-208_F | 5’ – AATGGATCCAAGAAGCACACAGGCTACGTCGGC – 3’ |
| USP7-559_R | 5’ – AATCTCGAGCTGCCGCTCCTTCCGCTTCTGAGC – 3’ |
| USP7-560_F | 5’ – AATGGATCCGAAGCCCATCTCTATATGCAAGTG – 3’ |
| USP7-800_R | 5’- AATCTCGAGATCACAGAAAATGACATCAACGCG – 3’ |
| USP7-801_F | 5’ – AATGGATCCAAAACAATCCCTAATGATCCTGGATTTGTG – 3’ |
| XIAP: BIR2+BIR3+UBA+RING-F | 5’ – GGCTTTTATCTTGAAAATAGTGCCAC – 3’ |
| XIAP_BIR2+BIR3+UBA+RING-R | 5’ – GCCAACTAAAACACTGCCATGTACATT – 3’ |
| XIAP_BIR1_F | 5’- AATGGATCCGAGAAGATGACTTTTAACAGTTTTG – 3’ |
| XIAP_BIR1_R | 5’- AATCTCGAGAGGGTTCCTCGGGTATATGG – 3’ |
| XIAP_BIR3+UBA+RING_F | 5’- AATCTCGAGAATATTCGAAGTGAATCTGATGCTG – 3’ |
| XIAP_BIR3+UBA+RING_R | 5’ – AATGCGGCCGCCAACTAAAACACTGCCATGTACATT – 3’ |
| XIAP_BIR1+BIR2_F | 5’- AATGGATCCGAGAAGATGACTTTTAACAGTTTTG – 3’ |
| XIAP_BIR1+BIR2_R | 5’- AATCTCGAGTGCCATGGATGGATTTCTTG – 3’ |
| XIAP_RING_F | 5’- AATCTCGAGTCACTTGAGGAGTGTCTGGT – 3’ |
| XIAP_RING_R | 5’ – AATGCGGCCGCCAACTAAAACACTGCCATGTACATT – 3’ |
| XIAP_BIR1+BIR2+BIR3+UBA_F | 5’- AATGGATCCGAGAAGATGACTTTTAACAGTTTTG – 3’ |
| XIAP_BIR1+BIR2+BIR3+UBA_R | 5’- AATGCGGCCGCCTAAAGCTTCTCCTCTTGCAGGC – 3’ |

#### *B. List of Real-Time PCR (RT-PCR) primers used*

| | |
|---|---|
| 18S rRNA_F | 5'- GCTTAATTTGACTCAACACGGGC – 3' |
| 18S rRNA_R | 5' – AGCTATCAATCTGTCAATCCTGTC – 3' |
| XIAP_F | 5' – TGGAATTTATGCTTTAGGTGAAGGT – 3' |
| XIAP_R | 5' – ACCCTGGATACCATTTAGCATGT – 3' |
| USP7_F | 5' – GCTCGACCACTTCAACAAAGC – 3' |
| USP7_R | 5' – CGTCCTCGCCTTGAACACA – 3' |
| Bcl-XL _ F | 5' – ACCCCAGGGACAGCATATCA – 3' |
| Bcl-XL_R | 5' – TGCGATCCGACTCACCAATA – 3' |
| Mdm2_F | 5' – GGCAGGGGAGAGTGATACAGA – 3' |
| Mdm2_R | 5' – GAAGCCAATTCTCACGAAGGG – 3' |
| p21_F | 5' – GCAGACCAGCATGACAGATTT – 3' |
| p21_R | 5' – GGATTAGGGCTTCCTCTTGGA – 3' |
| p53_F | 5' – GCTTTCCACGACGGTGAC – 3' |
| p53_R | 5' – GCTCGACGCTAGGATCTGAC – 3' |
| Bcl2_F | 5' – ATGTGTGTGGAGAGCGTCAACC – 3' |
| Bcl2_R | 5' – TGAGCAGAGTCTTCAGAGACAGCC – 3' |
| Bim_F | 5' – CTGCTGTCTCGATCCTCCAGT – 3' |
| Bim_R | 5' – GTCGTAAGATAACCATTCGTGGG – 3' |
| Bax_F | 5' – CTTTTGCTTCAGGGTTTCATC – 3' |
| Bax_R | 5' – TTGAGACACTCGCTCAGCTTC – 3' |
| Survivin_F | 5' – ACCACCGCATCTCTACATTCA – 3' |
| Survivin_R | 5' – CTCGTTCTCAGTGGGGCAGT – 3' |

### Supplementary Tables (S1-S4)

#### Table S1. USP7 interactome in HEK293T cells

| Uniprot acc No. | Gene Name | FC-A value |
|---|---|---|
| Q93009 | USP7 | 125.16 |
| P78347 | GTF2I | 74.16 |
| P09874 | PARP1 | 74.16 |
| P52272 | HNRNPM | 54.21 |
| P98170 | XIAP | 49.78 |
| P16152 | CBR1 | 40.91 |
| Q86U86 | PBRM1 | 40.91 |
| O75643 | SNRNP200 | 34.26 |
| Q7L2E3 | DHX30 | 32.04 |
| O00571 | DDX3X | 29.6 |
| Q9NZI8 | IGF2BP1 | 28.86 |
| Q8IX18 | DHX40 | 25.39 |
| P43243 | MATR3 | 25.39 |
| Q16643 | DBN1 | 23.17 |
| P42166 | TMPO | 23.17 |
| Q92922 | SMARCC1 | 23.17 |
| Q02218 | OGDH | 23.17 |
| Q6P2Q9 | PRPF8 | 23.17 |
| Q9BQG0 | MYBBP1A | 20.95 |
| Q15029 | EFTUD2 | 20.95 |
| Q8IX12 | CCAR1 | 20.95 |
| Q9NR30 | DDX21 | 20.95 |
| P11388 | TOP2A | 18.74 |
| Q92841 | DDX17 | 17.27 |
| Q9NYF8 | BCLAF1 | 16.94 |
| P08670 | VIM | 16.8 |
| Q8TDD1 | DDX54 | 16.52 |
| Q14498 | RBM39 | 16.52 |
| Q8TAQ2 | SMARCC2 | 16.52 |
| Q86YZ3 | HRNR | 16.52 |
| P36957 | DLST | 16.52 |
| P07910 | HNRNPC | 16.33 |
| Q9UHB6 | LIMA1 | 15.73 |
| P26368 | U2AF2 | 14.3 |
| Q9UMS4 | PRPF19 | 14.3 |
| Q9H0A0 | NAT10 | 14.3 |
| P46087 | NOP2 | 14.3 |

| | | |
|---|---|---|
| Q92878 | RAD50 | 14.3 |
| Q9BVP2 | GNL3 | 14.3 |
| Q9H307 | PNN | 14.3 |
| P17844 | DDX5 | 13.86 |
| Q9Y2W1 | THRAP3 | 12.29 |
| O76021 | RSL1D1 | 12.09 |
| Q99459 | CDC5L | 12.09 |
| Q9UQ35 | SRRM2 | 12.09 |
| Q9GZR7 | DDX24 | 12.09 |
| Q9P258 | RCC2 | 12.09 |
| P55265 | ADAR | 12.09 |
| O60264 | SMARCA5 | 12.09 |
| Q8IYB3 | SRRM1 | 12.09 |
| Q12906 | ILF3 | 12.09 |
| Q06787 | FMR1 | 12.09 |
| P23396 | RPS3 | 10.5 |
| Q8NC51 | SERBP1 | 10.28 |
| P02788 | LTF | 10.15 |
| Q6WCQ1 | MPRIP | 9.87 |
| P14373 | TRIM27 | 9.87 |
| P51114 | FXR1 | 9.87 |
| Q08170 | SRSF4 | 9.87 |
| P29692 | EEF1D | 9.87 |
| P62701 | RPS4X | 9.87 |
| Q9BZE4 | GTPBP4 | 9.87 |
| Q14247 | CTTN | 9.87 |
| Q9P0K7 | RAI14 | 9.87 |
| O95232 | LUC7L3 | 9.87 |
| Q01780 | EXOSC10 | 9.87 |
| O00425 | IGF2BP3 | 9.87 |
| Q6P0Q8 | MAST2 | 9.87 |
| O95793 | STAU1 | 9.87 |
| Q9BQ67 | GRWD1 | 9.87 |
| O43143 | DHX15 | 8.75 |
| Q08211 | DHX9 | 8.04 |
| A0A0A0MTS7 | TTN | 7.65 |
| P26640 | VARS | 7.65 |
| P20929 | NEB | 7.65 |
| A0A087WV90 | DMD | 7.65 |
| O75400 | PRPF40A | 7.65 |
| Q14669 | TRIP12 | 7.65 |
| Q7Z2W4 | ZC3HAV1 | 7.65 |
| E7ENN3 | SYNE1 | 7.65 |

| | | |
|---|---|---|
| Q09666 | AHNAK | 7.65 |
| Q9Y4I1 | MYO5A | 7.65 |
| P24534 | EEF1B2 | 7.65 |
| B2RWP5 | NSD1 | 7.65 |
| P08621 | SNRNP70 | 7.65 |
| Q15393 | SF3B3 | 7.65 |
| Q96GA3 | LTV1 | 7.65 |
| Q9NYQ8 | FAT2 | 7.65 |
| Q15149 | PLEC | 7.65 |
| P02768 | ALB | 7.55 |
| Q9BQ39 | DDX50 | 7.49 |
| P11940 | PABPC1 | 7 |
| O75533 | SF3B1 | 6.41 |
| P62424 | RPL7A | 6.2 |
| Q69YN4 | KIAA1429 | 5.43 |
| Q9HBM9 | MINK1 | 5.43 |
| O76027 | ANXA9 | 5.43 |
| Q59GJ9 | Acetyl-CoA carboxylase 2 variant | 5.43 |
| Q8IY21 | DDX60 | 5.43 |
| F8W705 | RGPD8 | 5.43 |
| E7ESB6 | GIGYF2 | 5.43 |
| E9PFH4 | TNPO3 | 5.43 |
| Q9UPN3 | MACF1 | 5.43 |
| C9IZF6 | PNPLA4 | 5.43 |
| Q7Z7G8 | VPS13B | 5.43 |
| H0YF42 | SLC38A6 | 5.43 |
| P35251 | RFC1 | 5.43 |
| F4MH73 | UTY | 5.43 |
| Q09028 | RBBP4 | 5.43 |
| Q13136 | PPFIA1 | 5.43 |
| P04637 | TP53 | 5.43 |
| Q9Y5B9 | SUPT16H | 5.43 |
| O14686 | KMT2D | 5.43 |
| Q9NX05 | FAM120C | 5.43 |
| Q9UKX3 | MYH13 | 5.43 |
| Q8TE73 | DNAH5 | 5.43 |
| Q86YA3 | C4orf21 | 5.43 |
| Q08E77 | UTP14C | 5.43 |
| P17480 | UBTF | 5.43 |
| K7ERG3 | TPM4 | 5.43 |
| Q58EX2 | SDK2 | 5.43 |
| E9PRE7 | PC | 5.43 |

| | | |
|---|---|---|
| B5LY64 | UMPS | 5.43 |
| O14746 | TERT | 5.43 |
| P01857 | IGHG1 | 5.43 |
| H7BZ39 | NGEF | 5.43 |
| Q07666 | KHDRBS1 | 5.43 |
| A4FU69 | EFCAB5 | 5.43 |
| B3KM90 | TPX2 | 5.43 |
| Q15751 | HERC1 | 5.43 |
| P67936 | TPM4 | 5.43 |
| Q9BY07 | SLC4A5 | 5.43 |
| B7WPL9 | PPIP5K1 | 5.43 |
| Q9BV73 | CEP250 | 5.43 |
| Q7RTY7 | OVCH1 | 5.43 |
| Q14789 | GOLGB1 | 5.43 |
| Q14676 | MDC1 | 5.43 |
| Q96T37 | RBM15 | 5.43 |
| B2R9K4 | Platelet-activating factor acetyl hydrolase | 5.43 |
| A6NGG8 | PCARE | 5.43 |
| Q96PE2 | ARHGEF17 | 5.43 |
| Q7Z6D5 | KLHL5 | 5.43 |
| Q02880 | TOP2B | 5.43 |
| Q6PIJ6 | FBXO38 | 5.43 |
| A0A0A0MR03 | CTDP1 | 5.43 |
| B4DFN2 | Dual specificity testis-specific protein kinase 2 | 5.43 |
| Q9H4Z2 | ZNF335 | 5.43 |
| Q96SB4 | SRPK1 | 5.43 |
| Q8N9T8 | KRI1 | 5.43 |
| Q99575 | POP1 | 5.43 |
| Q9NU22 | MDN1 | 5.43 |
| Q13435 | SF3B2 | 5.43 |
| P35968 | KDR | 5.43 |
| Q00610 | CLTC | 5.43 |
| Q9H1H9 | KIF13A | 5.43 |
| B7ZMN0 | LINGO2 | 5.43 |
| Q8IZT6 | ASPM | 5.43 |
| Q8NI27 | THOC2 | 5.43 |
| P55201 | BRPF1 | 5.43 |
| Q9NXJ9 | GON4L | 5.43 |
| D3DPG0 | TTN | 5.43 |
| Q15075 | EEA1 | 5.43 |
| B7ZLW1 | CAMSAP1 | 5.43 |

| | | |
|---|---|---|
| Q4LE36 | ACLY | 5.43 |
| Q96M86 | DNHD1 | 5.43 |
| Q5C9Z4 | NOM1 | 5.43 |
| Q0VF96 | CGNL1 | 5.43 |
| Q14432 | PDE3A | 5.43 |
| O15226 | NKRF | 5.43 |
| Q5TDG9 | DNAJC16 | 5.43 |
| Q86X02 | CDR2L | 5.43 |
| P42285 | SKIV2L2 | 5.43 |
| Q6PIF6 | MYO7B | 5.43 |
| Q6W2J9 | BCOR | 5.43 |
| H0YJV7 | YY1 | 5.43 |
| P12273 | PIP | 5.43 |
| P12270 | TPR | 5.43 |
| Q13610 | PWP1 | 5.43 |
| H0Y6V3 | CDC42BPA | 5.43 |
| O75475 | PSIP1 | 5.43 |
| Q9BQK8 | LPIN3 | 5.43 |
| Q15431 | SYCP1 | 5.43 |
| P49915 | GMPS | 5.43 |
| E9PEM5 | LRBA | 5.43 |
| Q8TDW7 | FAT3 | 5.43 |
| Q9P0U3 | SENP1 | 5.43 |
| Q9HC10 | OTOF | 5.43 |
| P25391 | LAMA1 | 5.43 |
| Q53FR4 | Vacuolar protein sorting 35 variant | 5.43 |
| Q05923 | DUSP2 | 5.43 |
| M1V481 | KIF5B-ALK_K17;A20 | 5.43 |
| Q53SF7 | COBLL1 | 5.43 |
| O43263 | ADARB1 | 5.43 |
| E9PDF6 | MYO1B | 5.43 |
| A0A087X0P0 | CENPE | 5.43 |
| H0Y9R5 | SNX25 | 5.43 |
| B2RP65 | CEP110 | 5.43 |
| H0YA13 | PRDM16 | 5.43 |
| P51532 | SMARCA4 | 5.43 |
| Q6GV26 | Macrophage oligopeptide transporter PEPT1 | 5.43 |
| Q9H0D6 | XRN2 | 5.43 |
| A0A087WYV5 | Slit homolog 2 protein | 5.43 |
| Q14980 | NUMA1 | 5.43 |
| Q2WGJ9 | FER1L6 | 5.43 |

| | | |
|---|---|---|
| K7EMN8 | NLRP11 | 5.43 |
| F5GYN0 | FAM186A | 5.43 |
| Q4KWH8 | PLCH1 | 5.43 |
| D3DSV6 | TEX15 | 5.43 |
| B4DR33 | highly similar to Ankyrin repeat domain-containing protein 32 | 5.43 |
| Q8IZL8 | PELP1 | 5.43 |
| H0YDE5 | KIAA1549L | 5.43 |
| F8W9J4 | Dystonin | 5.43 |
| Q4G138 | LEPR protein | 5.43 |
| B3KQF9 | highly similar to A-kinase anchor protein 9 | 5.43 |
| O14974 | PPP1R12A | 5.39 |
| P81605 | DCD | 5 |
| P38646 | HSPA9 | 4.93 |
| O00541 | PES1 | 4.85 |
| P14866 | HNRNPL | 4.77 |
| P62753 | RPS6 | 4.56 |
| Q5D862 | FLG2 | 4.39 |
| O43390 | HNRNPR | 4.35 |
| P05388 | RPLP0 | 4.21 |
| P15924 | DSP | 4.12 |
| P26641 | EEF1G | 4 |
| Q9UER7 | DAXX | 0.73 |

#### Table S2. XIAP interactome in HEK293T cells

| Uniport acc No. | Gene Name | FC-A Score |
|---|---|---|
| P98170 | XIAP | 26.07 |
| P0CG47 | UBB | 18.15 |
| Q8IW75 | SERPINA12 | 9.58 |
| O75369 | FLNB | 8.92 |
| P46781 | RPS9 | 7.76 |
| A0A087WWU8 | TPM3 | 5.62 |
| Q93009 | USP7 | 5.62 |
| Q9NR28 | DIABLO | 5.62 |
| P13639 | EEF2 | 4.96 |
| P08238 | HSP90AB1 | 4.96 |
| P27708 | CAD | 4.3 |
| B4DI39 | HS701 | 4.3 |
| Q9UHB6 | LIMA1 | 3.76 |
| Q7RTM4 | SYNE1 | 3.64 |
| O43464 | HTRA2 | 2.98 |
| Q0QF12 | SDHA | 2.98 |
| Q14686 | NCOA6 | 2.98 |
| P49411 | TUFM | 2.98 |
| P21333 | FLNA | 2.59 |
| Q96HS1 | PGAM5 | 2.32 |

#### Table S3. Ratio of proteome in P5091 vs DMSO treated HCT116 cells

| Uniprot ID | Ratio of P5091 vs DMSO | Protein name |
|---|---|---|
| P37108 | 1.51705665 | Signal recognition particle 14 kDa protein |
| P10412 | 1.448870814 | Histone H1.4 |
| P16402 | 1.448870814 | Histone H1.3 |
| P16403 | 1.448870814 | Histone H1.2 |
| P07919 | 1.370252789 | Cytochrome b-c1 complex subunit 6, mitochondrial |
| P99999 | 1.260153953 | Cytochrome c |
| O14737 | 1.215883343 | Programmed cell death protein 5 |
| O95881 | 1.151796529 | Thioredoxin domain-containing protein 12 |
| P14174 | 1.090712141 | Macrophage migration inhibitory factor |
| P62857 | 1.062597567 | 40S ribosomal protein S28 |
| P16949 | 1.023720196 | Stathmin |
| P39019 | 1.013148323 | 40S ribosomal protein S19 |
| O75347 | 0.998324656 | Tubulin-specific chaperone A |
| Q07955 | 0.992872995 | Serine/arginine-rich splicing factor 1 |
| P15311 | 0.980043842 | Ezrin |
| P09972 | 0.978610985 | Fructose-bisphosphate aldolase C |
| P09496 | 0.973193334 | Clathrin light chain A |
| P61604 | 0.955048746 | 10 kDa heat shock protein, mitochondrial |
| P18859 | 0.944083017 | ATP synthase-coupling factor 6, mitochondrial |
| P62273 | 0.940010242 | 40S ribosomal protein S29 |
| Q15056 | 0.923582352 | Eukaryotic translation initiation factor 4H |
| P06748 | 0.907063357 | Nucleophosmin |
| P68363 | 0.894735822 | Tubulin alpha-1B chain |
| Q71U36 | 0.894735822 | Tubulin alpha-1A chain |
| Q9BQE3 | 0.894735822 | Tubulin alpha-1C chain |
| P04792 | 0.876937532 | Heat shock protein beta-1 |
| P20962 | 0.870070779 | Parathymosin |
| Q86V81 | 0.865994836 | THO complex subunit 4 |
| P63241 | 0.861259025 | Eukaryotic translation initiation factor 5A-1 |
| P04075 | 0.860238756 | Fructose-bisphosphate aldolase A |
| P00441 | 0.854965967 | Superoxide dismutase [Cu-Zn] |
| P62805 | 0.853949936 | Histone H4 |
| Q13126 | 0.845857723 | S-methyl-5'-thioadenosine phosphorylase |
| Q92945 | 0.827206709 | Far upstream element-binding protein 2 |
| Q13263 | 0.822425128 | Transcription intermediary factor 1-beta |
| Q15847 | 0.811456962 | Adipogenesis regulatory factor |
| O14745 | 0.807469359 | Na(+)/H(+) exchange regulatory cofactor NHE-RF1 |
| P38646 | 0.804004037 | Stress-70 protein, mitochondrial |
| P43487 | 0.803023891 | Ran-specific GTPase-activating protein |
| P05787 | 0.775069526 | Keratin, type II cytoskeletal 8 |
| P25398 | 0.769890179 | 40S ribosomal protein S12 |
| P23588 | 0.769092162 | Eukaryotic translation initiation factor 4B |

| | | |
|---|---|---|
| P27816 | 0.762194458 | Microtubule-associated protein 4 |
| Q8N257 | 0.754103049 | Histone H2B type 3-B |
| O75531 | 0.750712332 | Barrier-to-autointegration factor |
| P10809 | 0.750579569 | 60 kDa heat shock protein, mitochondrial |
| Q99497 | 0.747183827 | Protein/nucleic acid deglycase DJ-1 |
| P62937 | 0.744897344 | Peptidyl-prolyl cis-trans isomerase A |
| O00193 | 0.73586171 | Small acidic protein |
| O75533 | 0.73155164 | Splicing factor 3B subunit 1 |
| O14979 | 0.721104036 | Heterogeneous nuclear ribonucleoprotein D-like |
| P11940 | 0.70004952 | Polyadenylate-binding protein 1 |
| P31948 | 0.694474073 | Stress-induced-phosphoprotein 1 |
| P05387 | 0.677481916 | Coiled-coil-helix-coiled-coil-helix domain-containing protein 10, mitochondria |
| Q8WYQ3 | 0.662153508 | Coiled-coil-helix-coiled-coil-helix domain-containing protein 10, mitochondria |
| Q13185 | 0.661836876 | Chromobox protein homolog 3 |
| Q99880 | 0.622487891 | Histone H2B type 1-L |
| P21333 | 0.617242526 | Filamin-A |
| P49773 | 0.590753515 | Histidine triad nucleotide-binding protein 1 |
| P60174 | 0.577844156 | Triosephosphate isomerase |
| P19338 | 0.559372084 | Nucleolin |
| P09651 | 0.556444726 | Heterogeneous nuclear ribonucleoprotein A1 |
| P98170 | 0.546841689 | E3 ubiquitin-protein ligase XIAP |
| Q96A08 | 0.53346064 | Histone H2B type 1-A |
| Q8WXI7 | 0.532006294 | Mucin-16 |
| Q9UK76 | 0.510686848 | Jupiter microtubule associated homolog 1 |
| P62633 | 0.463863269 | Cellular nucleic acid-binding protein |
| P04406 | 0.441519896 | Glyceraldehyde-3-phosphate dehydrogenase |
| Q96Q06 | 0.437216029 | Perilipin-4 |
| P06454 | 0.380079132 | Prothymosin alpha |
| P17096 | 0.32291661 | High mobility group protein HMG-I/HMG-Y |
| P02795 | 0.306591107 | Metallothionein-2 |
| P23527 | 0.297595859 | Histone H2B type 1-O |
| P33778 | 0.297595859 | Histone H2B type 1-B |
| P58876 | 0.297595859 | Histone H2B type 1-D |
| P62807 | 0.297595859 | Histone H2B type 1-C/E/F/G/I |
| Q16778 | 0.297595859 | Histone H2B type 2-E |
| Q5QNW6 | 0.297595859 | Histone H2B type 2-F |
| Q93079 | 0.297595859 | Histone H2B type 1-H |
| Q99877 | 0.297595859 | Histone H2B type 1-N |
| Q99879 | 0.297595859 | Histone H2B type 1-M |
| Q6DN03 | 0.082445083 | Putative histone H2B type 2-C |
| Q6DRA6 | 0.082445083 | Putative histone H2B type 2-D |
| O60814 | 0.062327498 | Histone H2B type 1-K |
| P06899 | 0.062327498 | Histone H2B type 1-J |
| P57053 | 0.059427304 | Histone H2B type F-S |

#### Table S4. Expressions of USP7 & XIAP in multiple cancer cell lines

**(Proteomics DB)**

| Cell Line | USP7 | XIAP |
|---|---|---|
| RPMI-8226 | 6.202146 | 4.366029 |
| HT55 | 6.123565 | 4.867517 |
| MOLT-4 | 6.109394 | 3.311396 |
| Hep-G2 | 6.060375 | 3.331262 |
| MFM-223 | 6.055049 | 4.255584 |
| HEK-293 | 6.027757 | 4.746657 |
| HCT-15 | 6.004715 | 3.178295 |
| CCRF-CEM | 5.98715 | 3.927211 |
| OXCO-1 | 5.968749 | 4.666907 |
| SF-295 | 5.944806 | 3.954476 |
| NCI-H460 | 5.944143 | 4.189964 |
| C80 | 5.924396 | 4.656301 |
| HDC-8 | 5.915825 | 4.931465 |
| RCM-1 | 5.915796 | 4.986507 |
| SW837 | 5.915598 | 4.529862 |
| C125-PM | 5.914561 | 4.859621 |
| RKO | 5.914464 | 4.52471 |
| UACC-62 | 5.888796 | 4.460832 |
| HCC-202 | 5.885205 | 4.152254 |
| HME Lonza | 5.868753 | 4.330151 |
| HCC-2998 | 5.867981 | 4.41331 |
| HDC-143 | 5.861374 | 4.891372 |
| SN-12C | 5.858329 | 4.551637 |
| HDC-114 | 5.857444 | 4.748857 |
| KM-12 | 5.850482 | 3.149778 |
| HeLa | 5.835316 | 4.914952 |
| NCI-H548 | 5.834032 | 4.519059 |
| CCK-81 | 5.831609 | 4.852471 |
| SW-620 | 5.807502 | 4.929947 |
| HMT-3522 | 5.803577 | 3.975941 |
| PC/JW | 5.802307 | 5.130301 |
| LNCaP | 5.789534 | 4.367418 |
| C70 | 5.78669 | 4.51 |
| Colo 320DM | 5.786304 | 4.475841 |
| HDC-9 | 5.76943 | 4.54532 |
| Colo-205 | 5.768534 | 4.190764 |
| LS1034 | 5.760517 | 4.711524 |
| GP2d | 5.750228 | 4.399981 |
| C99 | 5.738118 | 4.495139 |
| HCA-46 | 5.735318 | 4.936152 |
| SNU-C2B | 5.730783 | 4.749667 |
| C84 | 5.727473 | 4.700339 |

| | | |
|---|---|---|
| HDC-54 | 5.722659 | 4.336619 |
| SKOV-3 | 5.718048 | 4.03326 |
| HDC-142 | 5.714116 | 4.609091 |
| T-47D | 5.711078 | 3.622243 |
| HDC-111 | 5.695491 | 4.959203 |
| NCI-H23 | 5.689915 | 4.516208 |
| C32 | 5.686542 | 4.976693 |
| K-562 | 5.684642 | 4.182357 |
| HRA-19 | 5.683346 | 4.762202 |
| MCF-7 | 5.679232 | 4.230049 |
| NCI-H747 | 5.678819 | 5.142785 |
| U-251 MG | 5.677831 | 4.06665 |
| OVCAR-8 | 5.675829 | 3.826415 |
| SK-MEL-5 | 5.674012 | 4.350018 |
| MDA-MB-453 | 5.673034 | 3.638887 |
| C75 | 5.671053 | 4.979228 |
| HES-3 | 5.661012 | 3.598907 |
| JURKAT | 5.657435 | 4.785571 |
| PC-3 | 5.651743 | 3.579241 |
| SW-480 | 5.63551 | 4.877236 |
| HT-29 | 5.63355 | 4.400457 |
| HDC-57 | 5.629563 | 4.357927 |
| SK-N-BE (2) | 5.628436 | 3.840344 |
| OVCAR-5 | 5.619913 | 4.262234 |
| LS513 | 5.616382 | 4.74548 |
| SK-MEL-2 | 5.6045 | 4.69087 |
| SK-MEL-28 | 5.594654 | 3.400794 |
| VACO 4A | 5.586577 | 4.63028 |
| LS411 | 5.58437 | 4.800244 |
| OVCAR-8/ADR | 5.576212 | 4.22855 |
| CC07 | 5.572415 | 4.955003 |
| M14 | 5.551886 | 4.243351 |
| HCC-1937 | 5.549153 | 4.219631 |
| C106 | 5.534925 | 4.395653 |
| HDC-73 | 5.532827 | 4.754526 |
| SK-CO-1 | 5.531546 | 4.565247 |
| ACHN | 5.5304 | 4.261661 |
| Mixed | 5.529046 | 3.598907 |
| HDC-135 | 5.522487 | 4.379909 |
| LOX-IMVI | 5.519229 | 3.985445 |
| LS123 | 5.517355 | 4.580719 |
| 786O | 5.510455 | 4.397812 |
| OVCAR-4 | 5.479399 | 4.091742 |
| TK-10 | 5.466471 | 4.541791 |
| HDC-82 | 5.459048 | 3.836646 |
| HCC-1143 | 5.449482 | 4.545622 |
| A-498 | 5.446955 | 4.842205 |

| | | |
|---|---|---|
| GaMG | 5.446592 | 4.915635 |
| IGROV-1 | 5.440042 | 4.368359 |
| A-549 | 5.43797 | 5.009171 |
| RXF-393 | 5.4375 | 3.970573 |
| A-375 | 5.436443 | 4.002182 |
| HCC-2218 | 5.430013 | 3.766582 |
| HCC-1599 | 5.425696 | 3.309975 |
| HOP-62 | 5.425401 | 3.885808 |
| Colo 741 | 5.420244 | 4.706451 |
| CC20 | 5.411638 | 4.577813 |
| OXCO-3 | 5.401321 | 4.53648 |
| C10 | 5.386382 | 5.044088 |
| SF-268 | 5.373168 | 2.841163 |
| OVCAR-3 | 5.328122 | 4.304966 |
| SNB-19 | 5.307313 | 4.409577 |
| EKVX | 5.306298 | 4.350275 |
| SF-539 | 5.299923 | 4.492943 |
| NCI-H226 | 5.294619 | 3.793518 |
| Microglia/CD11b+ | 5.29458 | 3.299209 |
| MDA-MB-435 | 5.283253 | 4.553074 |
| HOP-92 | 5.276548 | 3.918593 |
| HMEpC | 5.255423 | 3.565184 |
| SNB-75 | 5.204183 | 4.390522 |
| Non-small cell lung cancer | 5.189998 | 3.239454 |
| Caki-1 | 5.088841 | 4.949809 |
| NCI-H322M | 5.016784 | 4.05387 |
| BT-549 | 5.014213 | 4.186506 |
| SR | 5.010794 | 4.006725 |
| Hs-578T | 4.949947 | 4.098982 |
| MDA-MB-231 | 4.916602 | 3.934339 |
| U2-OS | 4.851898 | 4.052274 |
